# Dopamine enhances model-free credit assignment through boosting of retrospective model-based inference

**DOI:** 10.1101/2021.01.15.426639

**Authors:** Lorenz Deserno, Rani Moran, Jochen Michely, Ying Lee, Peter Dayan, Raymond J. Dolan

## Abstract

Dopamine is implicated in signalling model-free (MF) reward prediction errors and various aspects of model-based (MB) credit assignment and choice. Recently, we showed that cooperative interactions between MB and MF systems include guidance of MF credit assignment by MB inference. Here, we used a double-blind, placebo-controlled, within-subjects design to test the hypothesis that enhancing dopamine levels, using levodopa, boosts the guidance of MF credit assignment by MB inference. We found that levodopa enhanced retrospective guidance of MF credit assignment by MB inference, without impacting on MF and MB influences per se. This drug effect positively correlated with working memory, but only in a context where reward needed to be recalled for MF credit assignment. The dopaminergic enhancement in MB-MF interactions correlated negatively with a dopamine-dependent change in MB credit assignment, possibly reflecting a potential trade-off between these two components of behavioural control. Thus, our findings demonstrate that dopamine boosts MB inference during guidance of MF learning, supported in part by working memory, but trading-off with a dopaminergic enhancement of MB credit assignment. The findings highlight a novel role for a DA influence on MB-MF interactions.

## Introduction

Dual system theories of reinforcement learning (RL) propose behaviour is controlled by competitive and cooperative interactions between a prospective, model-based (MB), planning system and a retrospective, model-free (MF), value-caching system (Daw and Dayan, 2014; Dolan and Dayan, 2013). MF value-caching is driven by reward prediction error (RPE) signalling via phasic dopamine (DA, Montague et al., 1996; Schultz et al., 1997; Steinberg et al., 2013), a finding mirrored in human neuroimaging studies (D’Ardenne et al., 2008; O’Doherty et al., 2004). While DA RPEs are assumed to train MF values (a process we refer to as MF credit assignment or MFCA), there is evidence that DA neuromodulation also impacts MB learning (MB credit assignment or MBCA) and control (Doll et al., 2012; Langdon et al., 2018). For example, the activity of DA neurons reflects MB values (Sadacca et al., 2016), DA RPEs reflect hidden-state inference (Starkweather et al., 2017), and optogenetic activation and silencing of DA neurons impact the efficacy of MB learning (Sharpe et al., 2017). Human studies also show that higher DA levels are linked to enhanced MB influences (Deserno et al., 2015; Doll et al., 2016; Sharp et al., 2016; Wunderlich et al., 2012), which was confirmed in a non-human animal study (Groman et al., 2019), potentially mediated by a modulation in the efficiency of working memory or motivation.

RL theory has proposed cooperative interactions between MB and MF systems, including the idea that a MB controller instructs a MF system about the structure of the environment (Daw and Dayan, 2014; Mattar and Daw, 2018; Sutton, 1991). For instance, inferences made in a MB manner can disambiguate different possible states of the world in cases in which the MF system is otherwise unable to learn properly because it does not know the state. We recently provided empirical evidence for this sort of MB-MF cooperation, showing that retrospective MB inference guides MFCA via provision of knowledge regarding the environment’s transition structure (Moran et al., 2019). Given DA’s contribution to both MF and MB systems, we set out to examine whether this aspect of MB-MF cooperation is subject to DA influence.

To address this question, we used a dual-outcome bandit task (Moran et al., 2019) in a double-blind, placebo-controlled, within-subjects pharmacological study, employing levodopa to boost the brain’s overall DA levels. This task allows a separate measurement of MB and MF systems, and specifically probes guidance of MF learning based on MB knowledge of the environmental transition structure. Our hypothesis was that enhancing DA would strengthen the guidance of MFCA by MB inference. Importantly, at the time MB inference is possible, some rewards are no longer perceptually available to participants. Thus, we expected that DA-induced boosting of the MB guidance of MFCA would depend on working memory capacity exclusively for perceptually absent rewards. Finally, in light of previous reports that levodopa enhanced MB influences (Sharp et al., 2016; Wunderlich et al., 2012), we examined whether this is also true for our dual-outcome task, expecting that inter-individual differences in the effect of boosting DA on MB influences and on MB guidance of MFCA would be related.

Foreshadowing our results, we found that boosting DA levels via levodopa enhanced guidance of a MFCA by MB inference, an effect moderated by inter-individual differences in working memory but only when reward needed to be recalled. While boosting DA did not alter the overall influence of a MB system on choice per se, the drug effects on guidance of MFCA by MB inference and on MB choice were negatively correlated.

## Results

### Study design and task logic

We conducted a placebo-controlled, double-blind, within-subjects pharmacological study using levodopa to enhance presynaptic DA levels, as in previous studies (Chowdhury et al., 2013; Wunderlich et al., 2012). Participants were tested twice, once under the influence of 150mg levodopa, and once on placebo, where drug order was counterbalanced across individuals (n=62, Figure 1A; cf. Methods). On each lab visit, participants performed a task first introduced previously by Moran et al. (2019). The task was framed as a treasure hunt game called the “Magic Castle”. Initially, participants were trained extensively on a transition structure between states, under a cover narrative of four vehicles and four destinations. Subjects learned that each vehicle (state) travelled to two different sequential destinations in a random order (Figure 1B). The mapping of vehicles and destinations remained stationary throughout a session, but the two test sessions featured different vehicles and destinations. At each destination, participants could potentially earn a reward with a probability that drifted across trials according to four independent random walks (Figure 1C).

**Figure 1.**
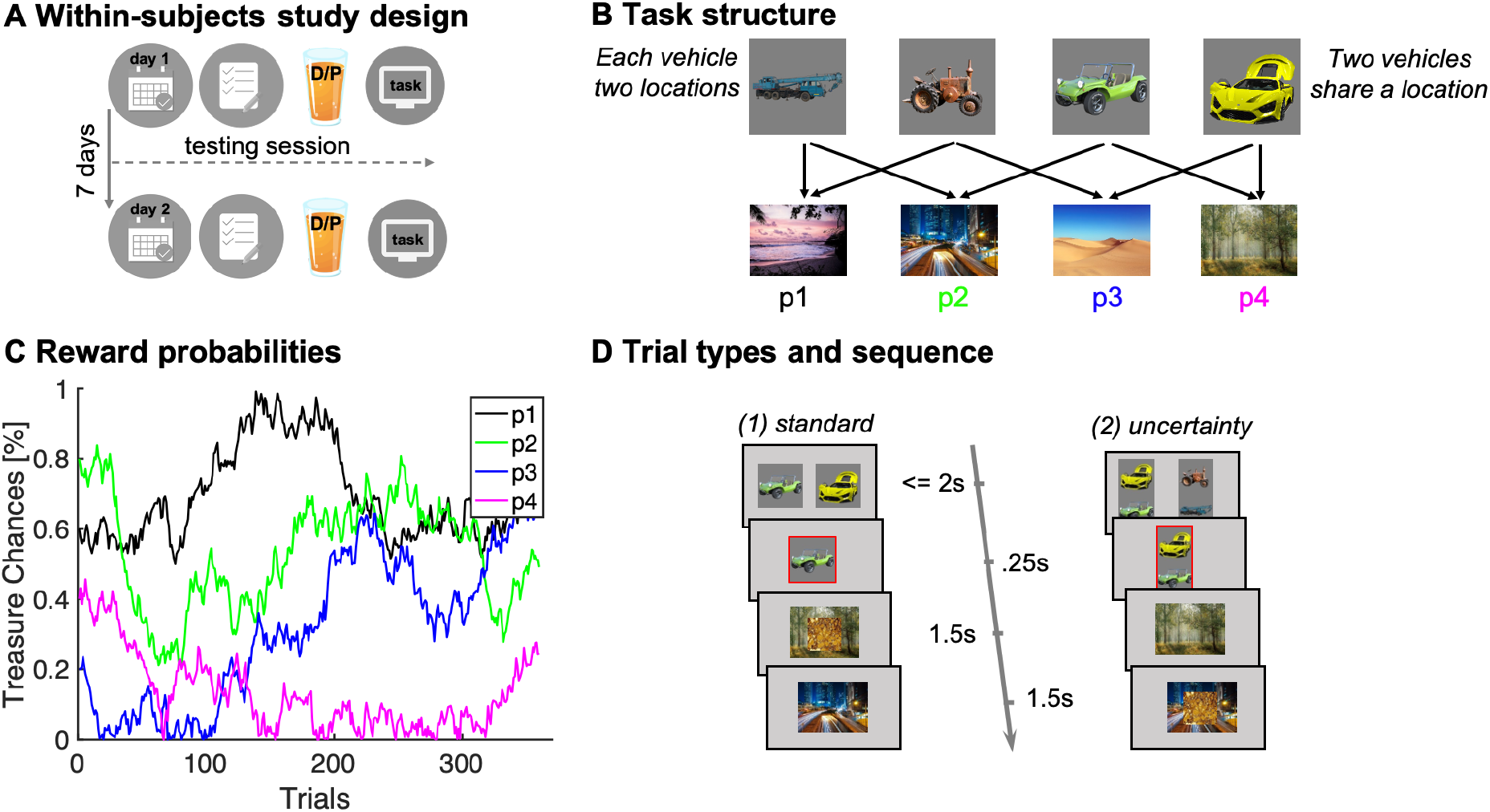
**A)** Illustration of within-subjects design. On each of two testing days, approximately 7 days apart, participants started with either a medical screening and brief physical exam (day 1) or a working memory test (day 2). Subsequently they drank an orange squash containing either levodopa (D) or placebo (P). **B)** Task structure of the Magic Castle Game. Following a choice of vehicle, participants “travelled” to two associated destinations. Each vehicle shared a destination with another vehicle. At each destination, participants could win a reward (10 pence) with a probability that drifted slowly as Gaussian random walks, illustrated in **C). D)** Depiction of trial types and sequences. (1) On *standard* trials (2/3 of the trials), participants made a choice out of two options in trial-n (max. choice 2s). The choice was then highlighted (.25s) and participants subsequently visited each destination (.5s displayed alone). Reward, if obtained, was overlaid to each of the destinations for 1s. (2) On *uncertainty trials*, participants made a choice between two pairs of vehicles. Subsequently, the ghost nominates, unbeknown to the participant, one vehicle out of the chosen pair. Firstly, the participant is presented the destination shared by the chosen pair of vehicles and this destination is therefore *non-informative* about the ghost’s nominee. Secondly, the destination unique to the ghost-nominated vehicle is then shown. This second destination is *informative* because it enables inference of the ghost’s nominee with perfect certainty based on a MB inference that relies on task transition structure. Trial timing was identical for standard and uncertainty trials.

The task included two trial types (Figure 1D): (1) standard trials (2/3 of the trials) and (2) uncertainty trials (1/3 of the trials). On standard trials, participants were offered two vehicles and upon choosing one, they visited both its associated destinations where they could earn rewards. On uncertainty trials, participants likewise chose a pair of vehicles (from two offered vehicle-pairs). Next, an unseen ghost randomly nominated a choice of one of the vehicles in the chosen pair, and a visit to its two destinations followed. Critically, participants were not privy to which vehicle was nominated by the ghost. However, they could resolve this uncertainty after seeing both visited destinations based on their knowledge of task transition structure. We refer to this as retrospective MB inference. Such inference can only occur after exposure to the second destination, as only then can subjects know which of the two vehicles the ghost had originally selected.

We first present ‘model-agnostic’ analyses focusing on how events on trial *n* affect choices on trial *n+1*. This allows identification of MF and MB choice signatures, the guidance of MFCA by retrospective MB inference, and, crucially, whether these signatures varied as a function of drug treatment (levodopa vs. placebo). These analyses are supported by validating simulations using computational models as provided in a later section.

### Logic of model-free and model-based contributions to choices

A MF system updates values based on earned rewards only for a chosen vehicle (illustrated in Figure 2A). A MB system does not maintain and update values for the vehicles directly. Instead, the MB system updates the values of destinations and calculates prospectively on-demand values for each offered vehicle (see computational modelling). This enables the MB system to generalize value across vehicles which share a common destination (illustrated in Figure 2B).

**Figure 2.**
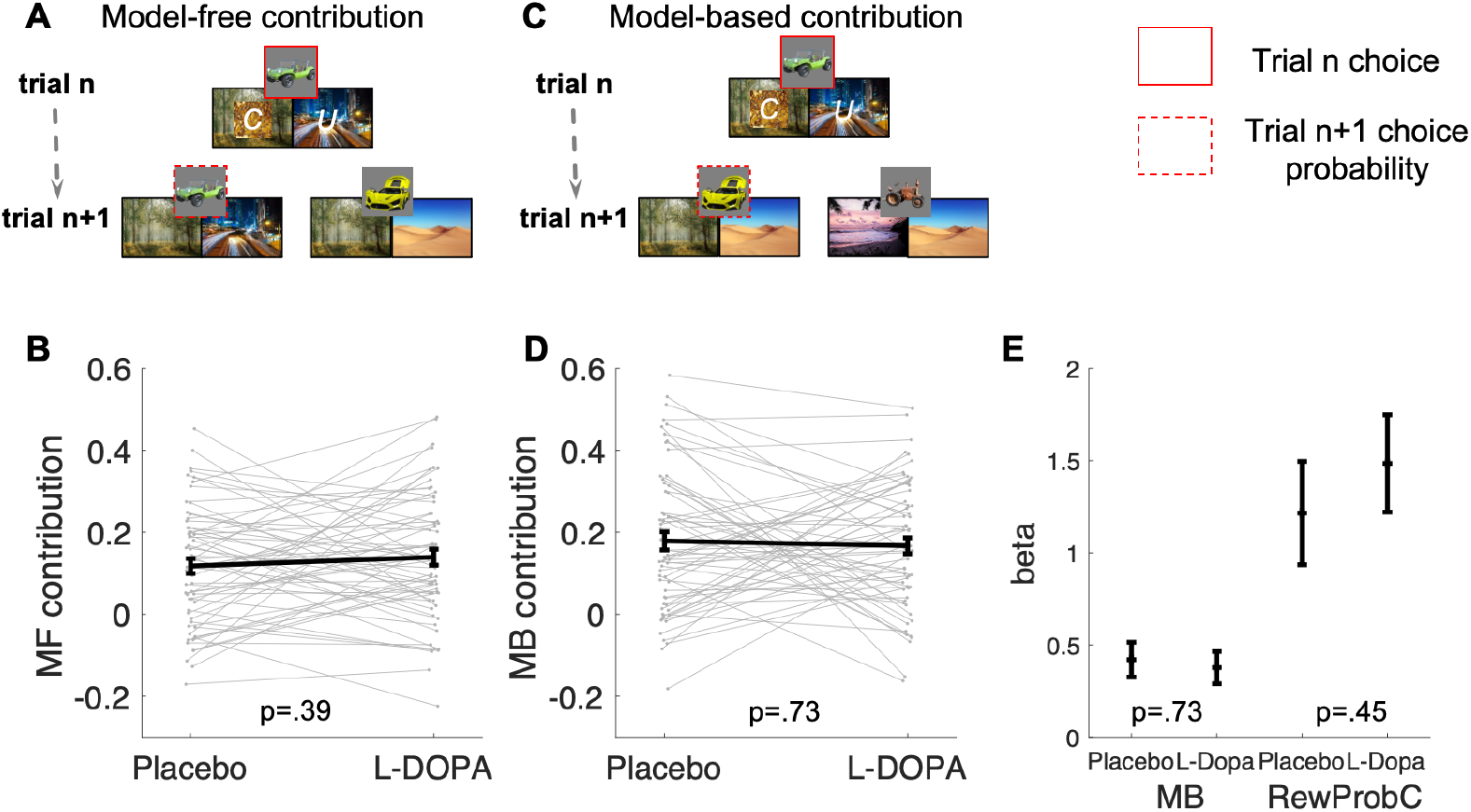
**A)** Illustration of MF choice repetition. We consider only standard trials n+1 that offer for choice the standard trial n chosen vehicle (e.g. green antique car) alongside another vehicle (e.g. yellow racing car), sharing a common destination. Following choice of a vehicle in trial n (framed in red), participants visited two destinations of which one can be labelled on trial n+1 as common to both offered vehicles (C, e.g. forest, which was also rewarded in the example) and the other labelled as unique (U, e.g. city highway, unrewarded in this example) to the trial n chosen vehicle. The trial n common-destination reward effect on the probability to repeat the previously chosen vehicle constitutes a MF choice repetition. **B)** The empirical reward effect at the common destination (i.e., the difference between rewarded and unrewarded on trial n, see Figure S3 for a more detailed plot) on repetition probability in trial n+1 is plotted for placebo and levodopa (L-DOPA) conditions. There was a positive common-reward main effect and this reward effect did not differ significantly between placebo and levodopa conditions. **C)** Illustration of the MB contribution. We considered only standard trials n+1 that excluded from the choice set the standard trial n chosen vehicle (e.g. green antique car). One of the vehicles offered on trial n+1 shared one destination in common with the trial-n chosen vehicle (e.g., yellow racing car and we term its choice a generalization). A reward (on trial n) effect for the common destination on the probability to generalize on trial n+1 constitutes a signature of MB choice generalization. **D)** The empirical reward effect at the common destination (i.e., the difference between rewarded and unrewarded, see Figure S3 for a more detailed plot) on generalization probability is plotted for placebo and levodopa conditions. **E)** In the regression analysis described in the text, we also include the current (subject- and trial-specific) state of the drifting reward probabilities (at the common destination) because we previously found this was necessary to control for temporal auto correlations in rewards (Moran et al., 2019). For completeness, we plot beta regression weights of reward versus no reward at the common destination (indicated as MB) and for the common reward probability (RewProbC) each for placebo and levodopa conditions. No significant interaction with drug session was observed. Error bars correspond to SEM reflecting variability between participants.

### No evidence of dopaminergic modulation for MF choice repetition

Consider a pair of standard trials n and n+1 for which the vehicle chosen on the former is also offered on the latter, against another vehicle (Figure 2A). The two vehicles offered on trial n+1 reach a common destination, but the vehicle previously chosen on trial n also visits a unique destination. In a logistic mixed effects model, we regressed a choice repetition of this vehicle on whether the common and/or unique destinations were rewarded on trial n (reward/non-reward) and on drug status (levodopa/placebo). Replicating a previous finding (Moran et al., 2019), we found a main effect for common reward (b=0.67, t(7251)=9.14, p<.001). This effect constitutes MF choice repetition, as the MB system appraises that the common destination favours both trial n+1 vehicles (see Figure S1 for validating simulations). As expected on both MB and MF grounds, there was a main effect for unique reward (b=1.54, t(7251)=17.40, p<.001). There was no drug × common-reward interaction (b=0.07, t(7251)=.67, p=.500), providing no evidence for a drug-induced change in MF choice repetition on standard trials (Figure, 2B). None of the remaining (main or interaction) effects were significant (Table S1).

### No evidence of dopaminergic modulation for MB choice generalization

Consider a standard trial-n+1, which excludes the vehicle chosen on trial n from the choice set. This trial-n chosen vehicle shares a destination with one of the trial-n+1 offered vehicles, allowing an analysis of MB choice generalization. Using a logistic mixed effects model, wherein we regressed choice generalization on trial-n rewards at the common destination, on the current reward probability of the common destination and on drug session, replicated our previous finding (Moran et al., 2019) of a positive main effect for the common-reward (b=0.40, t(7177)=6.22, p<.001). This positive common trial-n reward-effect on choice constitutes a MB choice generalization (even after controlling for the drifting reward probability at the common destination, see Figure S1 for validating simulations). The common-reward × drug interaction was not significant (b=0.05, t(7177)=0.39, p=.695), providing no evidence for a drug-induced change in MB choice (Figure 2D & E). Except for the main effect of the drifting reward probability at the common destination, no other effects were significant (Table S1).

In summary, we replicate previous findings (Moran et al., 2019) of mutual MF and MB contributions to choices. There was no evidence, however, that these contributions were modulated by levodopa.

### Retrospective MB inference guides MFCA

We next addressed our main question: Does levodopa administration boost a MB guidance of MFCA through a retrospective MB inference? In an uncertainty trial, participants choose one out of the two pairs of vehicles (Figure 1D). Next, a ghost randomly nominates a vehicle from the chosen pair (Figure 3). Participants then observe a destination common to both of the vehicles of the chosen pair, followed by a destination unique to the ghost-nominated vehicle. As participants are uninformed about the ghost nominee, they have a 50-50% belief initially and observing the first destination is *non-informative* with respect to the ghost’s nominee (as it is shared between vehicles). Critically, following observation of the second destination, a MB system can infer the ghost-nominated vehicle with absolute certainty based upon knowledge of the task transition structure. Thus, the second destination is retrospectively *informative* with respect to inference of the ghost’s nominee. Subsequently, the inferred vehicle information can be shared with a MF system to direct MFCA towards the ghost-nominated vehicle. We predicted guidance of MFCA occurs for both vehicles in the chosen pair, but to a different extent. Specifically, guidance of MFCA for the ghost-nominated, as compared to the ghost-rejected, vehicle would support an hypothesis that retrospective MB inference preferentially guides MFCA (Moran et al., 2019). See Figure S2 for validating model simulations. Our novel hypothesis here is that this effect will be strengthened under levodopa as compared to placebo, which we examine, firstly via the informative and, secondly, via the non-informative destination.

**Figure 3.**
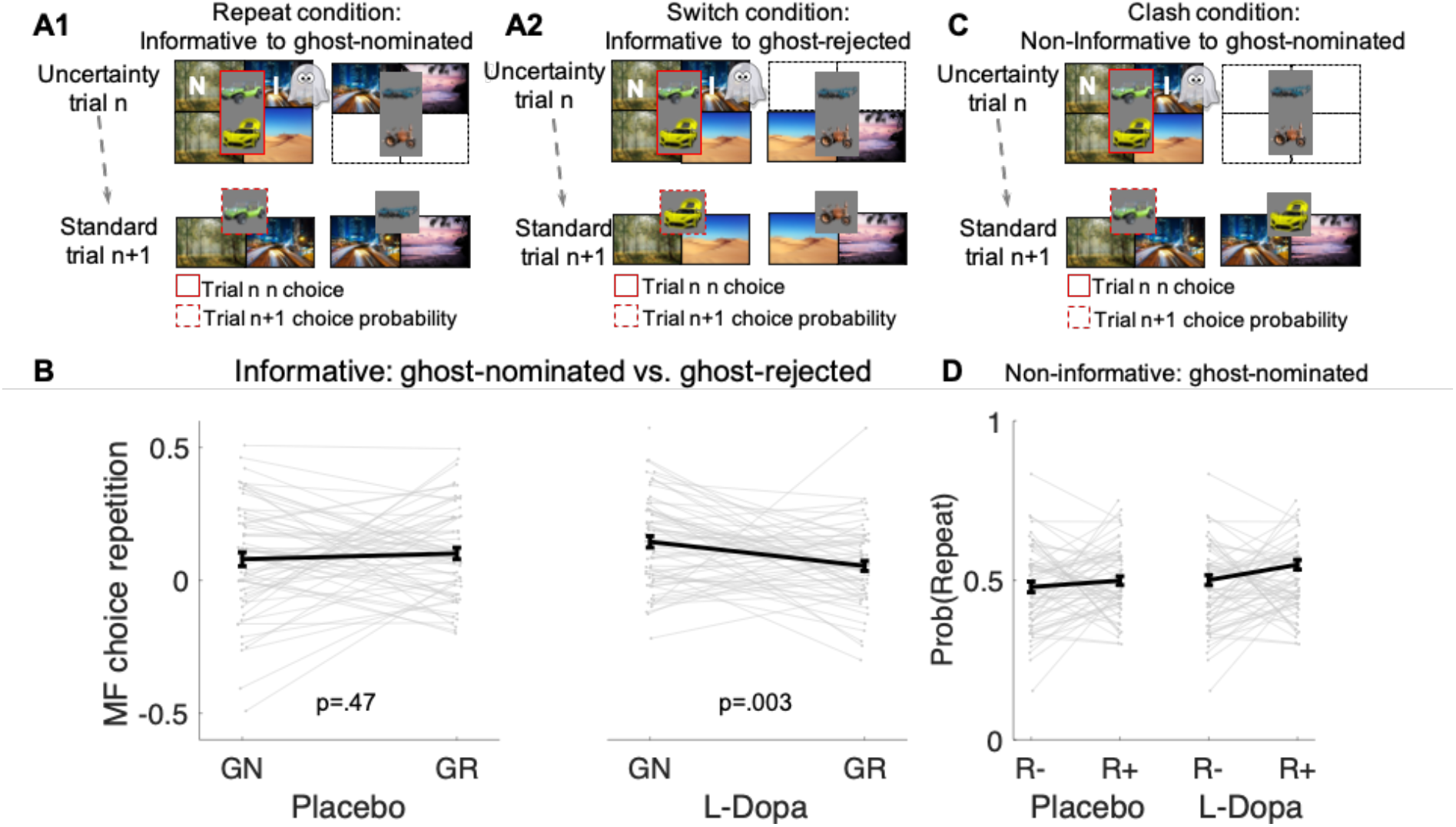
In an uncertainty trial n, participants choose a pair of vehicles. The ghost nominates one vehicle out of this pair (e.g., green antique car). Participants have a chance belief about the ghost-nominated vehicle. The firstly presented destination holds no information about the ghost-nominated vehicle, the non-informative (“N) destination. The destination presented second enables retrospective MB inference about the ghost’s nomination and is therefore informative (“I”). **A1.** Illustration of the repeat condition. The ghost-nominated vehicle (e.g., green antique car) is offered for choice in standard trial n+1 alongside a vehicle from the non-chosen pair (e.g., blue building crane). A higher probability to repeat the ghost-nominated vehicle in standard trial n+1 after a reward as compared to no reward at the informative destination constitutes MFCA for the ghost’s nomination (GN). **A2.** Illustration of the switch condition. The ghost-rejected vehicle (e.g., the yellow racing car) is offered for choice in standard trial n+1 alongside a vehicle from the non-chosen pair (e.g. brown farming tractor). A higher probability to choose the ghost-rejected vehicle in standard trial n+1 after a reward as compared to no reward at the informative destination constitutes MFCA for the ghost’s rejection (GR). Both ghost-based assignments depend on retrospective MB inference. **B.** Preferential effect of retrospective MB inference on MFCA (effects of GN>GR) based on the informative destination is enhanced under levodopa (L-Dopa) as compared to placebo. This is indicated by a significant trial type (GN/GR) × drug (placebo/ levodopa) interaction. Under levodopa, MFCA for GN is significantly higher than of GR, which is not the case under placebo (see Figure S4 for a more detailed plot). **C.** Illustration of the clash condition. The previously chosen pair is offered for choice in standard trial n+1. A higher probability to repeat the ghost-nominated vehicle in standard trial n+1 following reward (relative to non-reward) at the non-informative destination constitutes a signature of preferential MFCA for GN over GR. **D.** Choice repetition in clash trial is plotted as a function of reward and drug-group (see Figure S5 for a more detailed plot). While there was a main effect for drug, there was no interaction of non-informative reward × drug, providing no evidence that drug modulated MFCA based on the non-informative outcome. R+: reward; R-: non-reward. Error bars correspond to SEM reflecting variability between participants.

### Dopamine enhances preferential guidance of MFCA for the informative destination

MFCA for the ghost-nominated vehicle is tested in a “repeat” standard trial n+1 that follows an uncertainty trial n, as depicted in Figure 3 A1. MFCA of the ghost-rejected vehicle is examined in a “switch” standard trial n+1 following an uncertainty trial n, as depicted in Figure 3 A2. For a detailed analysis of repeat and switch trials, see Supplementary Information (SI) and Figure S4. The key metric of interest for our drug analysis is the contrast between MFCA for ghost-nominated *versus* ghost-rejected vehicles, based on the reward effects at the informative destination in repeat and switch trials (repeat or ghost-nominated / switch or ghost-rejected), separately for each nomination trial type (repeat/switch) × drug condition (levodopa /placebo) (Figure 3B). In a mixed effects model (Table S2), we found no main effect either of nomination (b=.043 t(239)=1.60, p=.110) or of drug (b=.01, t(239)=.40, p=.690). Crucially, we found a significant nomination × drug interaction (b=.11, t(239)=2.56, p=.011). A simple effects analysis revealed a preferential MFCA of the ghost-nominated over the ghost-rejected vehicle was significant under levodopa (b=.09, F(243,1)=9.07, p=.003) but not under placebo (b=-.02, F(243,1)=.53, p=.472). This supports our hypothesis that levodopa preferentially enhanced MFCA for the ghost-nominated, compared to ghost-rejected, vehicle under the guidance of retrospective MB inference. The nomination × drug interaction was not affected by session order (see Table S2).

### Dopaminergic modulation of preferential MFCA for the non-informative destination

A second means to examine MB influences over MFCA is to consider the non-informative destination. In a standard “clash” trial-n+1 following an uncertainty trial-n, the ghost-nominated vehicle is offered for choice alongside the ghost-rejected vehicle as depicted in Figure 3C. We previously showed that a positive effect of reward at the non-informative destination on choice repetition (i.e., a choice of the previously ghost-nominated vehicle) implicates a preferential guidance of MFCA towards the ghost-nominated vehicle guided by retrospective MB inference (Moran et al., 2019). In contrast, a MB system has knowledge that a non-informative destination is common to both standard trial n+1 vehicles. Note, this effect of reward at the non-informative destination can only occur when uncertainty about the ghost’s nomination was resolved retrospectively, once the informative destination was encountered.

In a logistic mixed effects model, we regressed choice repetition on trial-n rewards at informative and non-informative destinations as well as on drug session. A marginally significant main effect for the reward at the non-informative destination provides some support for preferential MFCA of the ghost-nominated vehicle (b=0.13, t(4861)=1.96, p=.051). Additionally, we found a main effect for reward at the informative destination (b=1.01, t(4861)=9.95, p<.001), as predicted by both the enhanced MFCA for the ghost-nominated vehicle and by an MB contribution. The interaction effect between drug and non-informative reward, however, was not significant (b=0.05, t(4861)=.39, p=.696, Figure 3D), nor were any other interactions in the model (Table S2). This analysis yielded no evidence that levodopa enhanced preferential guidance of MFCA based on reward at a non-informative destination. Unexpectedly, we found a positive main effect of drug (b=0.15, t(4861)=2.31, p=.021, Figure 3D), indicating that participants’ tendency to repeat choices of the ghost-nominated vehicle was generally enhanced under levodopa, but this finding that was only seen in this specific subset of trials and could not be corroborated based on computational modelling. We further dissect effects at the non-informative destination, in particular with respect to inter-individual differences in working memory, using computational modelling.

### Computational Modelling

One limitation of the analyses reported above is that they isolate the effects of the immediately preceding trial on a current choice. However, values and actions of RL agents are influenced by an entire task history and, to take account of such extended effects, we formulated a computational model that specified the likelihood of choices (Moran et al., 2019, also see Moran et al., in press, 2021). In brief, at choice, MF values (*Q*^MF^) of the two presented vehicles feed into a decision module. During learning, the MF system updates *Q*^MF^ of the *chosen* vehicle based on earned rewards alone. By contrast, the MB system prospectively calculates on-demand *Q*^MB^-values for each offered vehicle based on an arithmetic sum of the values of its two destinations:

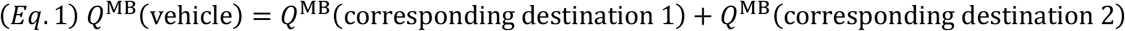

During learning, the MB system updates the values of the two visited destinations. We refer to these updates as MB credit assignment (MBCA). Unlike MFCA, which does not generalize credit from one vehicle to another, MBCA generalizes across the two vehicles which share a common destination. Thus, when a reward is collected in the forest destination, *Q*^MB^(forest) increases. As the forest is a shared destination, both vehicles that lead to this destination benefit during ensuing calculations of the on-demand *Q*^MB^-values. Critically, our model included five free “MFCA parameters” of focal interest, quantifying the extent of MFCA on standard trials (one parameter), on uncertainty trials (four parameters) for each of the objects in the chosen pair (nominated/rejected), and for each destination (informative/non-informative). We verified that the inclusion of these parameters was warranted using systematic model comparisons. A description of the sub-models and the model selection procedure is reported in the methods section and in Figure S6. We fitted our full model to each participant’s data in drug and placebo sessions based on Maximum Likelihood Estimation (see methods).

### Absence of dopaminergic modulation for MBCA and MFCA on standard trials

In line with our model-agnostic analyses of standard trials, we found positive contributions of MFCA (parameter 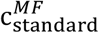; Fig. 4A) for both levodopa (M= 0.381, t(61)= 6.84, p<.001) and placebo (M= 0.326, t(61)= 5.76 p<.001), with no difference between drug conditions (t(61)= − 0.78, p=.442). Likewise, MBCA (parameter *c^MB^*; Fig. 4B) contributed positively for both levodopa (M= 0.255, t(61)= 7.88, p<.001) and placebo (M= 0.29, t(61)= 8.88, p<.001), with no significant difference between drugs (t(61)= 0.88, p=.3838). Thus, while both MBCA and MFCA contribute to choice, there was no evidence for a drug-related modulation. Forgetting and perseveration parameters of the model did not differ as a function of drug (see SI).

**Figure 4.**
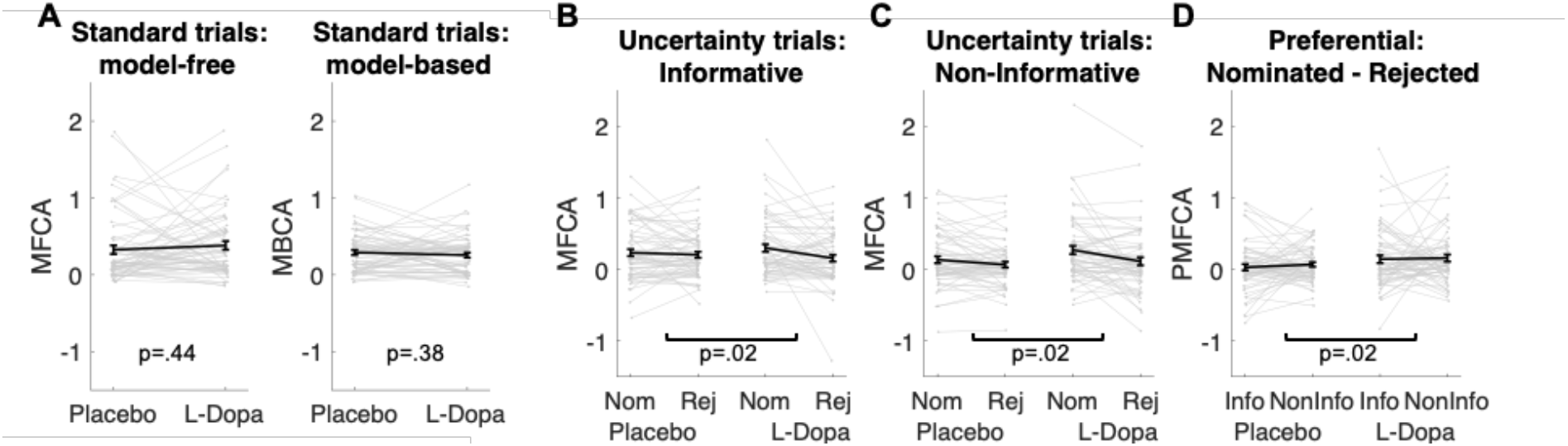
Analyses based on estimated credit assignment (CA) parameters from computational modelling. **A)** Model-free and model-based credit assignment parameters (MFCA; MBCA) did not differ significantly for placebo and levodopa conditions. **B)** MFCA parameters based on the informative outcome for the ghost-nominated and the ghost-rejected destinations as a function of drug condition. **D)** Same as C but for the non-informative destination. **E)** The extent to which MFCA prefers the nominated over the rejected vehicle for each destination and drug condition. We name this preferential MFCA (PMFCA).

### Levodopa enhances guidance of preferential MFCA by retrospective MB inference on uncertainty trials

To test our key hypothesis, that guidance of preferential MFCA by retrospective MB inference on uncertainty trials is enhanced by levodopa, we focused on the four computational parameters that pertaining to MFCA on uncertainty trials (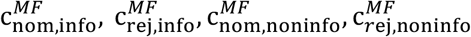. Figure 4B,C). In a mixed effects model we regressed these MFCA parameters on their underlying features: nomination (nominated / rejected), informativeness (informative / non-informative) and drug session (levodopa / placebo). Crucially, we found a positive nomination × drug interaction (b=0.10, t(480)=2.43, p=.015). A simple effects analysis revealed preferential MFCA (the effect of nomination) to be significant under levodopa (b=.13, F(488,1)=9.71, p=.002), and stronger than in the placebo condition (b=0.08, F(488,1)=4.83, p=.029), indicating that preferential MFCA was stronger under levodopa as compared to placebo. Importantly, this interaction was not qualified by a triple interaction (b=.02, t(480)=0.32, p=.738), providing no evidence that the extent of preferential MFCA differed for informative and non-informative outcomes. No other effect pertaining to drug reached significance (Table S3).

To examine in more fine-grained detail whether a MFCA is indeed preferential, we calculated, for each participant, in each session (drug/placebo), and for each level of informativeness (informative/non-informative), the extent to which MFCA was preferential for the ghost-nominated as opposed to the ghost-rejected vehicle (as quantified by 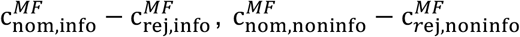; Figure 4D). Using a mixed effects model, we regressed preferential MFCA (PMFCA), based on MB guidance on informativeness and drug session.

We found a positive main effect for drug (b=0.10, t(240)= 2.41, p=.017), but neither the main effect of informativeness (b=-0.03, t(240)=-0.57, p=.568) nor the informativeness × drug interaction (b=.02, t(240)=0.33, p=.739) were significant. Using simple effects, MFCA preferred the ghost-nominated vehicle in the levodopa condition (b= 0.15, F(1,244)= 15.45, p<.001), while the same effect was only marginally significant in the placebo condition (b= 0.05, F(1,244)= 2.86, one-sided p=.046). Thus, our computational modelling analysis indicates that preferential MFCA is boosted by levodopa as compared to placebo across informative and non-informative destinations.

### Drug effect correlates positively with working memory only for reward at the non-informative destination

We hypothesized that working memory (WM) would moderate the boosting effect of levodopa, but only based on reward at the non-informative destination. When the informative destination is delivered on uncertainty trials, a MB system can infer the hidden choice and guide PMFCA. PMFCA based on reward at the non-informative destination can prefer the ghost-nominated vehicle only if it is at least partially postponed until uncertainty has been resolved by retrospective MB inference, in other words after delivery of the informative destination. At this time, reward received at the non-informative destination is no longer perceptually available and needs to be recalled (as illustrated in Figure 5A). Subjects’ WM capacity, as ascertained with the digit span test, showed a positive across-participants Spearman correlation with the drug effect (levodopa vs placebo) on PMFCA in the non-informative (r= .278, p=.029, Figure 5B), but not for the informative destination (r= −.057, p=.659, Figure 5B). The difference between these correlations was significant (p=. 044, permutation test; see methods). There was no significant correlation of WM capacity with drug-induced change in MBCA or with MBCA at levodopa or placebo (see SI).

**Figure 5.**
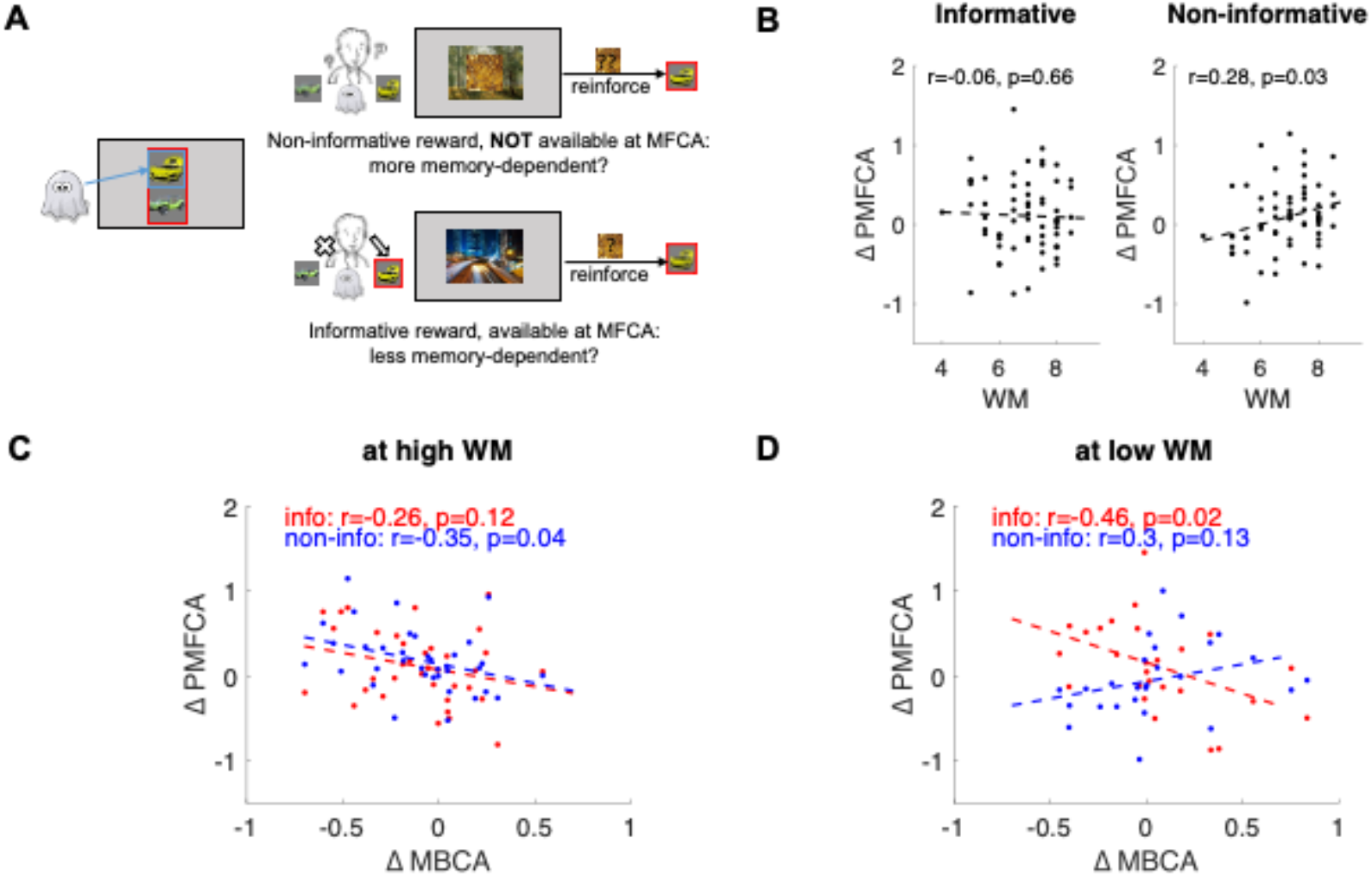
Inter-individual differences. **A)** Illustration of MFCA based on rewards at informative and non-informative destination. The latter is likely to depend more on memory recall because the reward is no longer perceptually available when MFCA can take place (after state uncertainty was resolved). **B)** Scatter plots of the drug effect (levodopa minus placebo) on preferential MFCA (Δ PMFCA) based on the informative destination reward and for the non-informative destination reward against working memory (WM). **C)** Scatter plot of the drug effect (levodopa minus placebo) on preferential MFCA (Δ PMFCA) based on the informative destination reward (info, red) and for the non-informative destination reward (non-info, blue) against drug-induced change in MBCA (Δ MBCA) at high working memory (WM) capacity. **D)** Same scatter plot as in C) but at low working memory (WM) capacity. In panels B, C and D regression lines are dashed. r refers to the Spearman correlation coefficient in panel B and Pearson correlation coefficient in C and D.

### Inter-individual differences in drug effects

Previous studies, using a task that cannot dissociate cooperative and competitive interactions between MB and MF systems, reported that boosting DA levels leads to enhanced MB choices (Sharp et al., 2016; Wunderlich et al., 2012), an effect we did not observe at a group level on our measure of MBCA. To explore the possibility that drug effects in different task conditions (guidance of MFCA vs. MBCA) are related, we analyzed inter-individual differences in the effects of boosting DA levels on guidance of MFCA and on MBCA. Because WM capacity correlated positively with drug effects at the non-informative destination as reported above, we included WM in the analysis of inter-individual differences in drug effects. Thus, we regressed DA-dependent differences (levodopa vs placebo) in PMFCA against informativeness, DA-dependent differences in MBCA and WM capacity. This model revealed an informativeness × MBCA × WM interaction (b=0.16, t(116)=2.16, p=.032). To unpack the interaction, we ran the model separately at high and low WM capacity based on a median split. In individuals with high WM capacity, this revealed a negative main effect of MBCA (b=-0.13, t(48)=-2.45, p=.018, see Figure 5C) which was not qualified by an interaction between informativeness × MBCA (b=-0.07, t(48)=-0.86, p=.40). This means that, for high WM individuals, the drug-effects on PMFCA and MBCA are negatively related for informative and non-informative destinations. In contrast, in individuals with low WM capacity, there was a significant negative informativeness × MBCA interaction (b=-0.23, t(68)=-2.43, p=.018; Figure 5D). A simple effects analysis revealed that the drug-effect on MBCA had a significant negative relation on the drug effect on PMFCA for the informative destination (b=-.18, F(1,68)=6.13, p=.015; Figure 5D) but not for the non-informative destination (b=.05, F(1,68)=0.42, p=.517; Figure 5D). Using model-agnostic metrics of DA-dependent change in guidance of MFCA and in MB choice, the negative correlation was also significant (see Figure S7). These inter-individual differences may reflect a trade-off between PMFCA and MBCA under boosted DA levels.

## Discussion

We show that enhancing dopamine boosted the guidance of model-free credit assignment by retrospective model-based inference. This pharmacological effect was associated with higher working memory capacity just for rewards that were no longer perceptually available and had to be recalled for credit assignment to be correct. Whereas both MF and MB influences were unaffected by the drug manipulation at the group level, analysis of inter-individual differences in drug effects showed that enhanced guidance of MFCA by retrospective MB inference was negatively correlated with drug-related change in MBCA. The findings provide, to our knowledge, the first human evidence that DA directly influences cooperative interactions between MB and MF systems, highlighting a novel role for DA in how MB information guides MFCA.

The effect of levodopa on prefrontal DA levels can lead to the enhancement of general aspects of cognition, for example WM (Cools and D’Esposito, 2011), probably depending on DA synthesis capacity in an inverted U-curved manner. The latter is likely to be important for supporting the computationally sophisticated operation of a MB system (Otto et al., 2013). One might therefore expect a primary drug effect on prefrontal DA to result in boosted MB influences (Sharpe et al., 2017; Wunderlich et al., 2012) – but we found no such influence. Equally, a long-standing proposal that phasic DA relates to a MF learning signal might predict that the main effect of the drug would be to speed or bias MF learning (Pessiglione et al., 2006). We observed no such effect, nor has it been seen in two previous studies (Sharp et al., 2016; Wunderlich et al., 2012). Instead, we found levodopa had a more specific influence, impacting the preferential MB guidance of MFCA in a situation where individuals needed to rely on retrospective MB inference to resolve state uncertainty. Thus, MB instruction about *what* (unobserved or inferred) state the MF system might *learn about*, was boosted under levodopa. In other words, DA boosts an exploitation of a model of task structure so as to facilitate retrospective learning about the past. These findings indicate an enhanced integration of MB information in DA signalling (Sadacca et al., 2016). Our results thus may provide a fine-grained view of the various processes involved – with the specificities of our task allowing us to separate out a rather particular component of WM, and an important, but restricted influence of MB information on MFCA.

First, preferential MFCA based on reward at the uninformative destination can only take place after seeing the informative destination and inferring the ghost’s choice. Thus, the uninformative destination’s reward has to be maintained in WM to support preferential MFCA. In other words, an ability to maintain information in working memory is a prerequisite for a DA-dependent boosting of preferential MFCA based on the uninformative destination. In line with this, we found a DA-boosting of MB guidance of MFCA depended on WM for the non-informative destination alone. This underlines the importance of accounting for inter-individual differences in supportive cognitive processes particularly when it comes to providing a detailed understanding of DA drug effects of interest (Cools, 2019; Kroemer et al., 2019).

Second, given that the information about the uninformative destination is stored in WM, what might be the neural mechanisms associated with its use in MB guidance of MFCA. Animal and human work points to a crucial role for orbitofrontal cortex in representing the model of a task model, including unobserved and inferred states, and in guiding behaviour accordingly (Howard et al., 2020; Jones et al., 2012; Schuck et al., 2016). This orbitofrontal function has also been related to the degree of sequential offline replay in the hippocampus (Schuck and Niv, 2019). Theoretical treatments of hippocampal offline neural replay proposes it informs credit assignment based on RPE (Mattar and Daw, 2018), a suggestion gaining support in recent empirical evidence in humans (Eldar et al., 2020; Liu et al., 2019, 2020). In our task, offline replay seems especially necessary to support preferential MFCA based on the first, uninformative, destination, because at this stage participants are still uncertain about the ghost’s choice. Under this account, we would predict enhanced offline replay (during rest between trials) of the non-informative destination (including its reward) and the inferred ghost’s choice under the influence of L-Dopa. Whether this enhanced replay occurs indirectly, via the interaction with WM, or is also a direct consequence of the L-Dopa is a pressing question for future work.

Previous studies, using a task not designed to test cooperative interactions between MB and MF systems (Daw et al., 2011), indicated a positive relationship between boosted DA and MB contributions to choice (Deserno et al., 2015; Doll et al., 2012; Sharp et al., 2016; Wunderlich et al., 2012). While MB choice contributions were not elevated at the group level by the drug in our data, we found a negative correlation between drug-related change on these contributions and on MB guidance of MFCA, in keeping with a trade-off between DA influences on these two components of behavioural control. In arbitrating between MB choice and retrospective MB inference to guide MFCA, participants need to weigh their respective cognitive costs vs. instrumental value. In independent recent work, a balance of costs and benefits was recently shown to be modulated by DA (Westbrook et al., 2020). Future studies will be needed to detail how the relative costs of planning vs. retrospective state-inference are influenced by DA, which can also inform DA contributions to trade-offs pertaining to strategy selection.

A limitation in our study is that guidance of informative MFCA by MB inference was significant in the levodopa condition alone but not in the placebo condition in model-agnostic measures (which are based on a subset of trials and consider only very recent influences on choice). However, computational modelling, informed by the entire trial-by-trial history of one’s experiences is arguably more sensitive, and this consideration enabled us to capture a preferential guidance of MFCA by MB inference also in the placebo condition.

In sum, our study provides first evidence that DA enhances cooperative interactions between MB and MF systems. The finding provides a unified perspective on previous research in humans and animals, suggesting a closely integrated architecture of how MF and MB systems interact under the guidance of DA-mediated so as to improve learning. DA-mediated cooperation between MB and MF control is a potentially exciting target for disentangling the precise role played by MB control in the development of impulsive and compulsive psychiatric symptoms.

## Methods

### Procedures

A total of 64 participants (32 females) completed a bandit at each of the two sessions with drug or placebo in counterbalanced order in a double-blinded design. One participant failed to reach required performance during training (see below) and task data could not be collected. Out of remaining 63 participants, one participant experienced side effects during task performance and was therefore excluded. Results reported above are based on a sample of n=62. All participants attended on two sessions approximately 1 week apart. Participants were screened to have no psychiatric or somatic condition, no regular intake of medication before invitation and received a short on-site medical screening at the beginning of their day 1 visit. At the beginning of the day 2 visit, they performed a working memory test, the digit span, which was thus only collected once.

### Drug protocol

The order of drug and placebo was counterbalanced. The protocol contained two decision-making tasks, which started at least 60min after ingestion of either levodopa (150 mg of levodopa + 37.5 mg of benserazide dispersed in orange squash) or placebo (orange squash alone with ascorbic acid). Benserazide reduces peripheral metabolism of levodopa, thus, leads to higher levels of DA in the brain and minimizes side effects such as nausea and vomiting. To achieve comparable drug absorption across individuals, subjects were instructed not to eat for up to 2h before commencing the study. Repeated physiological measurements (blood pressure and heart rate) and subjective mood rating scales were recorded under placebo and levodopa. A doctor prepared the orange squash such that data collection was double-blinded.

### Task Description

Participants were introduced to a minor variant of **a** task developed by Moran et al. (2019) using pictures of vehicles and destinations rather than objects and coloured rooms, and lasting slightly less time. The was presented as a treasure hunt called the ‘Magic Castle”. Before playing the main task, all participants were instructed that they can choose out of four vehicles from the Magic Castle’s garage that each vehicle could take them to two destinations (see Figure 1B). The mapping between vehicles and destination was randomly created for each participant and each session (sessions also had different sets of stimuli) but remained fixed for one session. They were then extensively trained on the specific vehicle-destination mapping. In this training, participants first saw a vehicle and had to press the space bar in self-paced time to subsequently visits the two associated destinations in random. The initial training run contained 12 repetitions per vehicle-destination mapping (48 trials). This training was followed by two types of each 8 quiz trials which asked to match one destination out of two to a vehicle or to match a vehicle out of two to a destination (time limit of 3sec). Each quiz trial had to be answered correctly and in time otherwise another training session was started with only 4 repetitions per vehicle-destination mapping (16 trials) followed again by the quiz. This procedure was repeated until participants passed all quiz. Participants were then introduced to the general structure of standard trials of bandit task (18 practice trials). This was followed by instructions introducing the ghost trials, which were complemented by another 16 practice trials including standard and ghost trials. Before starting the main experiment, participants performed a shorter refresher training of the vehicle-destination mapping with 4 repetitions per vehicle-destination mapping followed by the same quiz trials to passed as described above. In case of not passing at this stage, the refresher training was repeated with 2 repetitions per vehicle-destination mapping until the quiz was passed.

During the subsequent main task, participants should try to maximize their earnings. In each trial, they could probabilistically find a treasure (reward) at each of the two destinations (worth 1 penny). Reward probabilities varied over time independently for each of the four destinations according to Gaussian random walks with boundaries at p=0 and p=1 and a standard deviation of .025 per trial (Figure 1C). Random walks were generated anew per participant and session. A total of 360 trials split in 5 blocks of each 72 trials were played with short enforced breaks between blocks. Two of three trials were ‘standard trials’, in which a random pair of objects was offered for choice sharing one common outcome (choice time <= 2s). After making a choice, they visited each destination subsequently in random order. Each destination was presented for 1s and overlaid with treasure or not (indicating a reward or not). The lag between the logged choice and the first destination as well as between first and second destinations was 500ms. Every third trial was an “uncertainty trial” in which two disjoint pairs of vehicles were offered for choice. Crucially, each of the presented pairs of vehicles shared one common outcome. Participants were told before the main task that after their choice of a pair of vehicles, the ghost of the Magic Castle would randomly pick one vehicle out of the chosen pair. Because this ghost was transparent, participants could not see the ghost’s choice. However, participants visited the two destinations subsequently and collected treasure reward (or not). Essentially, when the ghost nominated a vehicle, the common destination was presented first and the destination unique to this vehicle was presented second. At this time of presentation of the unique destination, participants could retrospectively infer the choice made by the ghost. Trial timing was identical for standard and ghost trials. The 120 standard trials following a previous trial n-1 standard trial included 30 presentations of each of the four eligible pairs of vehicles in a random order. The 120 uncertainty trials included 60 presentations of the two eligible pairings in a random order. The standard trials following uncertainty trials were defined according to the observed transition based on the (ghost’s) choice in the preceding (uncertainty) trial. These 120 trials contained 40 presentations of each of the “repeat”, “switch” or “clash” trial types in a random order. A repeat trial presented the ghost-nominated object alongside its vertical counterpart, a switch trial presented the ghost-rejected object alongside its vertical counterpart and a clash trial presented the previously selected pair.

### Model-agnostic analysis

Model agnostic analyses were performed with logistic mixed effects models using MATLAB’s “fitglme” function with participants serving as random effects with a free covariance matrix. All models included the variable ORDER as regressor (coded as +.5 for the first and −.5 for the second session) to control for unspecific effects and participants (PART) served as random effects. Details of are reported in Table S1.

The analysis of MF and MB contributions is restricted to standard trials followed by a standard trial. For MF contributions, we consider only a trial-n+1, which offers the trial-n chosen object for choice (against another object). Regressors C (common destination) and U (unique destination) indicated whether rewards were received at trial n (coded as +.5 for reward and −.5 for no reward) and were included to predict the variable REPEAT indicating whether the previously chosen vehicle was repeated or not. The variable DRUG was included as regressor indicating within-subject Levodopa or placebo session (coded as +.5 for levopdopa and −.5 for placebo). The model, in Wilkinson notation, can be found on Table S1. For MB contributions, we specifically examined trials in which the trial-n chosen vehicle was excluded on trial n+1. The regressors C, PART and DRUG were coded as for the analysis of the MF contribution. One additional regressor P was included, which coded the reward probability of the common destination and was centralized by subtracting .5. These regressors were included to predict the variable GENERALIZE indicated whether the choice on trial n+1 was generalized (choosing the vehicle not shown in trial n+1 that shares a destination with the trial-n chosen vehicle). The model, in Wilkinson notation, can be found on Table S1.

The analysis of how retrospective MB inference preferentially guides MFCA focused on standard trials following uncertainty trials. The key analysis reported above focuses on MF choice repetition for the ghost-nominated in contrast to the ghost-rejected vehicle. This was achieved by extracting empirical choice proportions from “repeat trials” and from “switch trials”. More specifically, we computed the proportion of repeating or switching after a reward minus no reward at the informative destination averaged across rewards at the non-informative destination (reflecting the main effect of the informative destination, “I”) for each trial type. These two metrics were subjected to a mixed-effects models as dependent variable and with TYPE (nominated / rejected coded as +.5 and −.5) and, as before, DRUG and PART as predictors. The model, in Wilkinson notation, can be found on Table S2. A detailed analysis using separate mixed effects models for repeat and switch conditions is reported in the SI.

Another model-agnostic analysis examined learning for the ghost-nominated and – rejected vehicles based on the uncertainty trial n non-informative destination and therefore focused on n+1 “clash” trials, which offer for choice the same pair of objects as chosen on the previous uncertainty trial (the ghost-nominated and ghost-rejected objects). Choice repetition was defined as choice of the ghost-nominated vehicle from uncertainty trial n indicated by the variable REPEAT. Regressors PART, N, I and DRUG are coded as previously. The model, in Wilkinson notation, can be found on Table S2.

### Computational Models

We formulated a hybrid RL model to account for the series of choices for each participant. In the model, choices are contributed by both the MB and MF systems. The MF system caches a *Q*^MF^-value for each vehicle, subsequently retrieved when the vehicle is offered for choice. During learning on standard trials, following reward-feedback, rewards from the two visited destinations are used to update the *Q*^MF^-value for the chosen vehicle as follows:

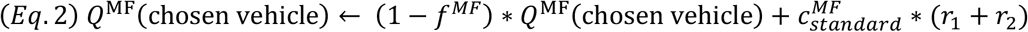

where 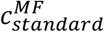 is a free MFCA parameter on standard trials and the r’s are the rewards for each of the two obtained outcomes (coded as 1 for reward or −1 for non-reward) and *f^MF^* (between 0-1) is a free parameter corresponding to forgetting in the MF system.

During learning on uncertainty trials, the MF values of the ghost nominated and ghost rejected options were updated according to:

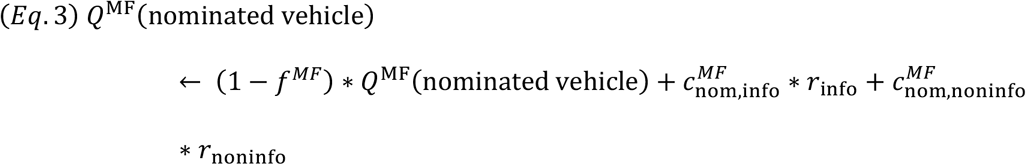

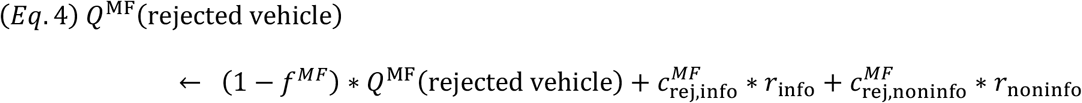

Where the c’s are free MFCA parameters on uncertainty trials for each destination (informative/non-informative) and vehicle type (ghost nominated/rejected) in the chosen pair. The r’s are rewards (once more, coded as 1 or −1) for the informative and non-informative outcomes.

The MF values of the remaining vehicles (3 on standard trials; 2 on uncertainty trials) were subject to forgetting:

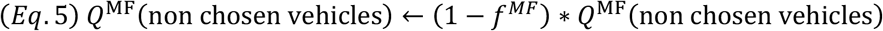

Unlike MF, the MB system maintains *Q*^MB^-values for the four different destinations. During choices the *Q*^MB^-value for each offered vehicle is calculated based on the transition structure (i.e., the two destinations associated with a vehicle):

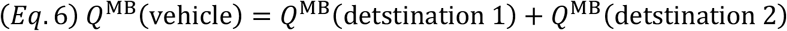

Following a choice (on both standard and uncertainty trials), the MB system updates the *Q*^MB^-values of each of the two observed destination based on its own reward:

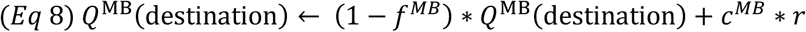

Where *f*^MB^ (bet. 0-1) is a free parameter corresponding to forgetting in the MB system, *c^MB^* is a free MBCA parameter and r corresponds to the reward (1 or −1) obtained at the destination.

Our model additionally included progressive perseveration for vehicles. After each standard trial the perseveration values of each of the 4 vehicles updated according to

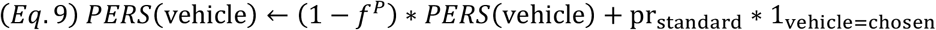

Where 1_vehicle=chosen_ is the chosen vehicle indicator, pr_standard_ is a free perseveration parameter for standard trials, and *f^P^* (bet. 0-1) is a free perseveration forgetting parameter. Similarly after each uncertainty trials perseverations values were updated according to:

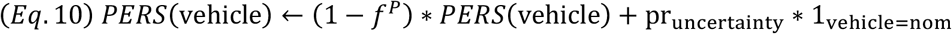

where 1_vehicle=nom_ is the ghost-nominated vehicle indicator, and pr_uncertainty_ is a free perseveration parameter for uncertainty trials.

During a standard trial choice a net Q value was calculated for each offered vehicle:

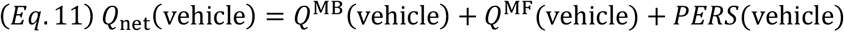

Similarly, during an uncertainty-trial choice the *Q*_net_ value of each offered vehicle-pair was calculated as a sum of the MB, MF and PERS values of that pair. MF, MB, and PERS values for a vehicle-pair in turn were each calculated as the corresponding average value of the two vehicles in that pair. For example:

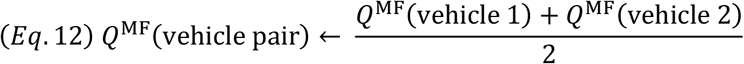

The *Q*_net_ values for the 2 vehicles offered for choice on standard trials are then injected into a softmax choice rule such that the probability to choose an option is:

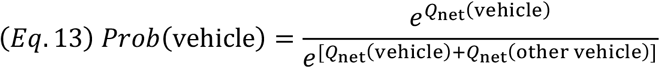

Similarly, on uncertainty trials the probability to choice a vehicle pair was based on softmaxing the net Q-values of the two offered pairs. *Q*^MF^ and *PERS* person-values and *Q*^MB^ vegetables-values where initialized to 0 at the beginning of the experiment.

### Model Comparison and Fitting

Our full hybrid agents, which allowed for contributions from both an MB and an MF system, served as a super-model in a family of six nested submodels of interest: 1) a pure MB model, which was obtained by setting the contribution of the MF to 0 (i.e. 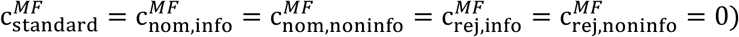), 2) a pure MF-action model, which was obtained by setting the contribution of the MB system to choices to 0 (i.e. *c^MB^* = 0; Note that in this model, MB inference was still allowed to guide MF inference), 3) a ‘no informativeness effect on MFCA’ sub-model obtained by constraining equality between the MFCA for the informative and non-informative destination (i.e., 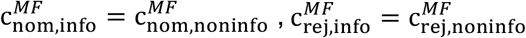), 4) a ‘no MB guided MFCA’ sub-model obtained by constraining equality between the MFCA parameters, for both the informative and non-informative destination, for the ghost-nominated and rejected objects 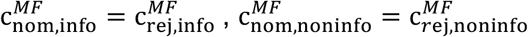, 5) a ‘no MB guidance of MFCA for the informative outcome’ obtained by constraining equality between the MFCA parameters for the ghost-nominated and ghost-rejected objects for the informative outcome 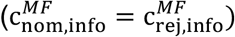, and 6) a ‘no MB guidance of MFCA for the non-informative outcome’ which was similar to 5 but for the non-informative outcome 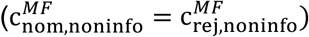.

We conducted a bootstrapped generalized likelihood ratio test, BGLRT (Moran and Goshen-Gottstein, 2015), for the super-model vs. each of the sub-models separately. In a nutshell, this method is based on the classical-statistics hypothesis testing approach and specifically on the generalized-likelihood ratio test (GLRT). However, whereas GLRT assumes asymptotic Chi-squared null distribution for the log-likelihood improvement of a super model over a sub-model, in BGLRT these distributions are derived empirically based on a parametric bootstrap method. In each of our model comparison the sub model serves as the H0 null hypothesis whereas the full model as the alternative H1 hypothesis. For each participant and drug condition, we created 1001 synthetic experimental sessions by simulating the sub-agent with the ML parameters on novel trial sequences which were generated as in the actual data. We next fitted both the super-agent and the sub-agent to each synthetic dataset and calculated the improvement in twice the logarithm of the likelihood for the full model. For each participant and drug condition, these 1001 likelihood-improvement values served as a null distribution to reject the sub-model. The p-value for each participant in each drug condition was calculated based on the proportion of synthetic dataset for which the twice logarithm of the likelihood-improvement was at least as large as the empirical improvement. Additionally, we performed the model comparison at the group level. We repeated the following 10,000 times. For each participant and drug condition we chose randomly, and uniformly, one of his/her 1,000 synthetic twice log-likelihood super-model improvements and we summed across participant and drug conditions. These 10,000 obtained values constitute the distribution of group super-model likelihood improvement under the null hypothesis that a sub-model imposes. We then calculated the p-value for rejecting the sub-agent at the group level as the proportion of synthetic datasets for which the super-agent twice logarithm of the likelihood improvement was larger or equal to the empirical improvement in super-model, summed across participants. Results, as display in Figure S6 in detail, fully supported the use of our full model including all effects of interest regarding MFCA in uncertainty trials.

We next fit our choice models to the data of each individual, separately for each drug condition (levodopa/placebo) maximizing the likelihood (ML) of their choices (we optimized likelihood using MATLAB’s ‘fmincon’, with 200 random starting points per participant * drug condition; Table S4 for distribution best-fitting parameters). See Table S2 for the distribution of full model’s fitted parameters.

### Model simulations

To generate model predictions with respect to choices, we simulated for each participant and each drug condition, 25 synthetic experimental sessions (novel trial sequences were generated as in the actual experiment), based on ML parameters obtained from the corresponding model fits. We then analysed these data in the same way as the original empirical data (but with datasets that were 25 times larger, as compared to the empirical data, per participant). Results are reported in Figures S1 and S2 of the SI. We also tested recoverability of model parameters (see Figure S7).

### Analysis of model parameters

All models included the variable ORDER as regressor (coded as +.5 for the first and −.5 for the second session) to control for unspecific effects and participants (PART) served as random effects. Details of are reported in Table S2.

For each participant in each drug condition, we obtained, based on the full model, four MFCA parameter estimates corresponding to destination (informative/non-informative) and vehicle (nominated/rejected) types. We conducted a mixed effects model (again implemented with MATLAB’s function “fitglme”) with TYPE (nominated/rejected coded as +.5 and −.5), INFO (informative/non-informative coded as +.5 and −.5) and DRUG (drug/placebo coded as +.5 and −.5) as regressors. The model, in Wilkinson notation, can be found on Table S3.

After finding significant drug by NOM * DRUG interaction, we followed this up in detail: we calculated for each participant in each drug condition and for each destination type the “preferential MFCA” (denoted PMFCA) effect as the difference between the corresponding nominated and rejected MFCA parameters. We next ran a mixed effects model for PMFCA. Our regressors where the destination type (denoted INFO; coded as before), and DRUG (coded as before). The model, in Wilkinson notation, can be found on Table S3.

### Correlations between preferential MFCA and WM

For each participant and destination type (Informative/non-informative), we contrasted the “preferential MFCA” estimates (as defined in the previous section) for levodopa minus placebo to obtain a drug-induced PMFCA effect. For each destination, we calculated across-participants Spearman correlations between these drug induced effects and WM. We compared the two correlations (for informative and non-informative destinations) using a permutation test. First, we z-scored the PMFCA separately for each destination type. Next we repeated the following steps (1-3), 10,000 times: 1) For each participant we randomly reshuffled (independent of other participants) the outcome type labels “informative” and “non-informative”, 2) We calculated the “synthetic” Spearman correlations between drug induced PMFCA effects and WM for each outcome type subject to the relabelling scheme and, 3) We subtracted the two correlations (non-informative minus informative). These 10,000 correlation-differences constituted a null distribution for testing the null hypothesis that the two correlations are equal. Finally, we calculated the p-value for testing the hypothesis of a stronger correlation for the non-informative destination as the percentage (of the 10,000) synthetic correlation-differences that were at least as large (in absolute value) as the empirical correlation-difference.

### Relationship between drug effects

We used the same score for drug-dependent change in PMFCA (levodopa minus placebo) and regressed it against informativeness, drug-dependent change in MBCA and working memory capacity in a mixed effects model.

## Acknowledgements

RJD is supported by a Wellcome Trust Investigator Award (098362/Z/12/Z) under which the above study was carried out. This work was carried out whilst R.J.D. was in receipt of a Lundbeck Visiting Professorship (R290-2018-2804) to the Danish Research Centre for Magnetic Resonance. RM is supported by the Max Panck Society and LD was at the time when the study was performed. The UCL-Max Planck Centre for Computational Psychiatry and Ageing is funded by a joint initiative between UCL and the Max Planck Society. RJD and LD are supported by a grant from the German Research Foundation (DFG TRR 265, project A02) and YL was at the time when the study was performed. For the purpose of Open Access, the authors have applied a CC BY public copyright license to any Author Accepted Manuscript version arising from this submission.

## Supplementary Information

### Repeat and switch standard trials following uncertainty trials

We showed previously on “repeat” trials (Moran et al., 2019), a positive effect of an informative destination reward (on trial n) on choice-repetition implicates MFCA to the ghost-nominated object (while the MB system knows that the value of the informative destination favours both vehicles on trial n+1). We also ran a separate analysis that examined MFCA for the ghost-nominated alone. In trial n+1 “repeat” trials, the ghost-nominated vehicle from trial-n is offered for choice alongside a vehicle from the trial-n non-chosen pair that shares the inference-allowing destination with the ghost-nominated object. Choice repetition was defined as choice of the ghost-nominated vehicle from uncertainty trial n as indicated by the variable REPEAT. Regressor PART is coded as previously. Regressors N (non-informative destination) and I (informative destination) indicate whether a reward was received at the destinations or not in trial n (coded as +.5/-.5). The model is REPEAT ~ N*I + (N*I | PART). This showed a main effect for the informative (I) destination (b=0.60, t(4885)=7.56, p=4e-14), supporting MFCA to the ghost-nominated object. Additionally, we found a main effect for the non-informative (N) destination (b=1.23, t(4885)=10.83, p=9e-42) as predicted by both MF and by MB contributions, and a significant interaction between the Informative and Non-informative destinations (b=0.31, t(4885)=2.09, p=.04). See Figure S4, A & B.

We showed previously on switch trials that a positive main effect of the informative outcome reward on choice-switching implicates MFCA for the ghost-rejected vehicle (because the MB system knows the informative destination is unrelated to both vehicles on trial n+1). A second separate analysis examined MFCA for the ghost-rejected vehicle In uncertainty trialn+1 “switch” trials, the ghost-rejected vehicle from trial-n is offered for choice alongside a vehicle from the trial-n non-chosen pair that shares a destination with the ghost-rejected object. Choice switching was defined as choice of the ghost-rejected vehicle from uncertainty trial n as indicated by the variable SWITCH. Regressors PART, N and I are coded as previously. The model is SWITCH ~ N*I + (N*I | PART). This showed a main effect for the reward at informative destination (b=0.38, t(4866)=5.60, p=2e-8), supporting MFCA to the ghost-rejected vehicle. While, this challenges any notion of *perfect* MB guidance of MFCA, it is consistent with the possibility that some participants, at least some of the time, do not rely on MB-inference because when MB inference does not occur, or when it fails to guide MF credit-assignment, the MF system has no basis to assign credit unequally to both vehicles in the selected pair. Additionally, we found a main effect for the non-informative destination reward (b=0.98, t(4866)=13.18, p=5e-39), as predicted by an MF credit-assignment to the ghost-rejected vehicle account but also by MB contributions. We found no significant interaction between rewards at the informative and non-informative destinations (Table S1). In Figure S2, we plot empirical choice proportions from both repeat and switch conditions (reflecting the effects reported above) in the manner as in the original paper by Moran et al. (2019) but separately for drug and placebo conditions. See Figure S4 C & D.

### Absence of drug effects on perseveration and forgetting parameters of the computational model

No difference between drug conditions was observed for perseveration parameter on standard trials (t(61)= 0.48, p=.63), perseveration parameter on uncertainty trials (t(61)= 0.51, p=.61), MF forgetting parameter (t(61)= 1.37, p=.17), MB forgetting parameter (t(61)= −0.33, p=.74), perseveration forgetting parameter (t(61)=0.30, p=.77).

### Correlation between WM and MBCA

Working memory moderated the boosting drug effect on guidance of MFCA based on retrospective MB inference but only based on non-informative reward (see main text). No moderating effect of working memory on a drug-dependent difference in MBCA was observed (r=-.07, p=.59). Working memory correlated positively with MBCA separately at placebo and at drug but this was non-significant (placebo: r=.21, p=.08; drug: r=.15, p=.23).

**Table S1.** Mixed-effects models on model-agnostic choice data from standard trials

| Name | Estimate | SE | tStat | DF | pValue | LowerCI | UpperCI |
| --- | --- | --- | --- | --- | --- | --- | --- |
| **MF choice (standard trials)** |  |  |  |  |  |  |  |
| REPEAT ~ 1+ C*U*DRUG*ORDER + (C+U+DRUG+ORDER \| PART) |  |  |  |  |  |  |  |
| (Intercept) | 0.34 | 0.06 | 5.54 | 7251 | .000 | 0.22 | 0.46 |
| C (common) | 0.67 | 0.07 | 9.14 | 7251 | .000 | 0.53 | 0.81 |
| U (unique) | 1.54 | 0.09 | 17.40 | 7251 | .000 | 1.36 | 1.71 |
| DRUG | 0.03 | 0.07 | 0.46 | 7251 | .643 | -0.11 | 0.18 |
| ORDER | 0.07 | 0.07 | 0.91 | 7251 | .365 | -0.08 | 0.21 |
| C*U | 0.19 | 0.11 | 1.72 | 7251 | .085 | -0.03 | 0.40 |
| C*DRUG | 0.07 | 0.11 | 0.67 | 7251 | .500 | -0.14 | 0.29 |
| U*DRUG | 0.06 | 0.11 | 0.56 | 7251 | .577 | -0.15 | 0.27 |
| C*ORDER | 0.12 | 0.11 | 1.09 | 7251 | .276 | -0.09 | 0.33 |
| U*ORDER | -0.11 | 0.11 | -0.99 | 7251 | .321 | -0.32 | 0.11 |
| DRUG*ORDER | -0.25 | 0.24 | -1.02 | 7251 | .309 | -0.73 | 0.23 |
| C*U*DRUG | 0.14 | 0.22 | 0.64 | 7251 | .524 | -0.29 | 0.57 |
| C*U*ORDER | 0.13 | 0.22 | 0.59 | 7251 | .554 | -0.30 | 0.56 |
| C*DRUG*ORDER | -0.02 | 0.29 | -0.06 | 7251 | .952 | -0.59 | 0.56 |
| U*DRUG*ORDER | -0.18 | 0.35 | -0.51 | 7251 | .609 | -0.87 | 0.51 |
| C*U*DRUG*ORDER | -0.22 | 0.44 | -0.50 | 7251 | .618 | -1.07 | 0.64 |
| **MB choice (standard trials)** |  |  |  |  |  |  |  |
| GENERALIZE ~ C*P*DRUG*ORDER + (C+P+DRUG+ORDER \| PART) |  |  |  |  |  |  |  |
| (Intercept) | 0.30 | 0.04 | 6.96 | 7177 | .000 | 0.22 | 0.38 |
| C (common) | 0.40 | 0.06 | 6.22 | 7177 | .000 | 0.27 | 0.52 |
| P (common reward probability) | 1.33 | 0.21 | 6.39 | 7177 | .000 | 0.92 | 1.74 |
| DRUG | -0.13 | 0.08 | -1.65 | 7177 | .099 | -0.29 | 0.03 |
| ORDER | -0.13 | 0.08 | -1.57 | 7177 | .116 | -0.29 | 0.03 |
| C*P | -0.23 | 0.23 | -1.01 | 7177 | .311 | -0.67 | 0.21 |
| C*DRUG | 0.05 | 0.12 | 0.39 | 7177 | .695 | -0.19 | 0.28 |
| P*DRUG | -0.34 | 0.23 | -1.48 | 7177 | .140 | -0.79 | 0.11 |
| C*ORDER | -0.06 | 0.12 | -0.52 | 7177 | .606 | -0.30 | 0.17 |
| P*ORDER | 0.16 | 0.23 | 0.70 | 7177 | .482 | -0.29 | 0.61 |
| DRUG*ORDER | -0.24 | 0.17 | -1.41 | 7177 | .158 | -0.58 | 0.09 |
| C*P*DRUG | -0.08 | 0.45 | -0.18 | 7177 | .856 | -0.97 | 0.80 |
| C*P*ORDER | 0.57 | 0.45 | 1.26 | 7177 | .207 | -0.31 | 1.45 |
| C*DRUG*ORDER | -0.38 | 0.25 | -1.48 | 7177 | .140 | -0.87 | 0.12 |
| P*DRUG*ORDER | 0.46 | 0.83 | 0.55 | 7177 | .583 | -1.18 | 2.09 |
| C*P*DRUG*ORDER | 1.40 | 0.91 | 1.54 | 7177 | .123 | -0.38 | 3.17 |

**Table S2.** Mixed-effects models on model-agnostic choice data from uncertainty trials

| Name | Estimate | SE | tStat | DF | pValue | LowerCI | UpperCI |
| --- | --- | --- | --- | --- | --- | --- | --- |
| **Preferential MFCA for the informative destination (Ghost-nominated, “repeat trials” > ghost-rejected, “switch trials”)** |  |  |  |  |  |  |  |
| MFCA ~ NOM*DRUG*ORDER + (NOM*DRUG+ORDER \| PART) |  |  |  |  |  |  |  |
| (Intercept) | 0.10 | 0.01 | 8.54 | 239 | .000 | 0.07 | 0.12 |
| NOM (Nomination) | 0.03 | 0.02 | 1.60 | 239 | .110 | -0.01 | 0.08 |
| DRUG | 0.01 | 0.02 | 0.40 | 239 | .690 | -0.04 | 0.05 |
| ORDER | 0.00 | 0.02 | 0.15 | 239 | .878 | -0.04 | 0.05 |
| NOM*DRUG | 0.11 | 0.04 | 2.56 | 239 | .011 | 0.03 | 0.20 |
| NOM*ORDER | 0.02 | 0.04 | 0.45 | 239 | .650 | -0.07 | 0.11 |
| DRUG*ORDER | -0.03 | 0.04 | -0.74 | 239 | .463 | -0.12 | 0.06 |
| NOM*DRUG*ORDER | -0.01 | 0.09 | -0.06 | 239 | .951 | -0.18 | 0.16 |
| **MFCA for non-informative destination (Ghost-nominated > ghost-rejected, “clash trials”)** |  |  |  |  |  |  |  |
| REPEAT ~ N*I*DRUG*ORDER + (N*I*DRUG+ORDER \| PART) |  |  |  |  |  |  |  |
| (Intercept) | 0.05 | 0.04 | 1.27 | 4861 | .203 | -0.03 | 0.12 |
| N (non-informative) | 0.13 | 0.07 | 1.96 | 4861 | .051 | 0.00 | 0.26 |
| I (informative) | 1.01 | 0.10 | 9.95 | 4861 | .000 | 0.81 | 1.21 |
| DRUG | 0.15 | 0.07 | 2.31 | 4861 | .021 | 0.02 | 0.29 |
| ORDER | 0.03 | 0.07 | 0.41 | 4861 | .684 | -0.10 | 0.16 |
| N*I | 0.08 | 0.14 | 0.57 | 4861 | .568 | -0.19 | 0.35 |
| N*DRUG | 0.05 | 0.13 | 0.39 | 4861 | .696 | -0.21 | 0.31 |
| I*DRUG | 0.03 | 0.14 | 0.24 | 4861 | .810 | -0.25 | 0.32 |
| N*ORDER | -0.05 | 0.13 | -0.34 | 4861 | .733 | -0.30 | 0.21 |
| I*ORDER | 0.06 | 0.14 | 0.43 | 4861 | .664 | -0.22 | 0.35 |
| DRUG*ORDER | -0.20 | 0.15 | -1.37 | 4861 | .171 | -0.49 | 0.09 |
| N*I*DRUG | 0.07 | 0.29 | 0.26 | 4861 | .798 | -0.49 | 0.64 |
| N*I*ORDER | 0.25 | 0.29 | 0.86 | 4861 | .388 | -0.32 | 0.81 |
| N*DRUG*ORDER | -0.47 | 0.26 | -1.80 | 4861 | .072 | -0.99 | 0.04 |
| I*DRUG*ORDER | -0.12 | 0.41 | -0.31 | 4861 | .759 | -0.92 | 0.67 |
| N*I*DRUG*ORDER | 0.86 | 0.55 | 1.56 | 4861 | .118 | -0.22 | 1.94 |

**Table S3.** Mixed-effects models on parameters of the computational model.

| Name | Estimate | SE | tStat | DF | pValue | LowerCI | UpperCI |
| --- | --- | --- | --- | --- | --- | --- | --- |
| **MFCA for ghost-nominated vs. ghost-rejected and informative vs non-informative**<br>MFCA ~ NOM*INFO*DRUG* + (NOM*INFO*DRUG \| PART) |  |  |  |  |  |  |  |
| (Intercept) | 0.18 | 0.02 | 7.60 | 480 | .000 | 0.14 | 0.23 |
| NOM (nomination) | 0.10 | 0.03 | 3.72 | 480 | .000 | 0.05 | 0.15 |
| INFO (informativeness) | 0.08 | 0.04 | 2.19 | 480 | .029 | 0.01 | 0.15 |
| DRUG | 0.05 | 0.05 | 1.00 | 480 | .316 | -0.05 | 0.15 |
| ORDER | 0.04 | 0.05 | 0.78 | 480 | .434 | -0.06 | 0.14 |
| NOM*INFO | -0.03 | 0.05 | -0.57 | 480 | .567 | -0.12 | 0.06 |
| NOM*DRUG | 0.10 | 0.04 | 2.43 | 480 | .015 | 0.02 | 0.18 |
| INFO*DRUG | -0.08 | 0.07 | -1.16 | 480 | .247 | -0.22 | 0.06 |
| NOM*ORDER | 0.02 | 0.04 | 0.37 | 480 | .715 | -0.07 | 0.10 |
| INFO*ORDER | 0.10 | 0.07 | 1.42 | 480 | .157 | -0.04 | 0.23 |
| DRUG:ORDER | -0.09 | 0.10 | -0.98 | 480 | .328 | -0.28 | 0.10 |
| NOM*INFO*DRUG | 0.02 | 0.07 | 0.33 | 480 | .738 | -0.12 | 0.17 |
| NOM*INFO*ORDER | -0.01 | 0.07 | -0.08 | 480 | .934 | -0.15 | 0.14 |
| NOM*DRUG*ORDER | -0.06 | 0.11 | -0.60 | 480 | .551 | -0.27 | 0.15 |
| INFO*DRUG*ORDER | 0.16 | 0.14 | 1.10 | 480 | .272 | -0.12 | 0.44 |
| NOM*INFO*DRUG*ORDER | 0.10 | 0.19 | 0.55 | 480 | .585 | -0.26 | 0.47 |
| **Preferential MFCA for informative vs. non-informative**<br>PMFCA ~ INFO*DRUG*ORDER + (INFO*DRUG+ORDER \| PART) |  |  |  |  |  |  |  |
| (Intercept) | 0.10 | 0.03 | 3.72 | 240 | .000 | 0.05 | 0.15 |
| INFO (informativeness) | -0.03 | 0.05 | -0.57 | 240 | .568 | -0.12 | 0.07 |
| DRUG | 0.10 | 0.04 | 2.41 | 240 | .017 | 0.02 | 0.18 |
| ORDER | 0.02 | 0.04 | 0.36 | 240 | .717 | -0.07 | 0.10 |
| INFO*DRUG | 0.02 | 0.07 | 0.33 | 240 | .739 | -0.12 | 0.17 |
| INFO*ORDER | -0.01 | 0.07 | -0.08 | 240 | .934 | -0.15 | 0.14 |
| DRUG*ORDER | -0.06 | 0.11 | -0.60 | 240 | .551 | -0.27 | 0.15 |
| INFO*DRUG*ORDER | 0.10 | 0.19 | 0.55 | 240 | .585 | -0.27 | 0.47 |

**Table S4.** Distribution of parameters from the full computational model.

| Cond. | % | MFCA standard | MFCA info-nom | MFCA info-rej | MFCA non-info-nom | MFCA non-info-rej | MBCA | perseveration - standard | perseveration - nominated | forget_MF | forget_MB | forget_Pers |
| --- | --- | --- | --- | --- | --- | --- | --- | --- | --- | --- | --- | --- |
| Placebo | 25 | 0.053 | -0.056 | -0.026 | -0.070 | -0.074 | 0.059 | -0.197 | -0.093 | 0.002 | 0.038 | 0.010 |
|  | 50 | 0.147 | 0.168 | 0.149 | 0.048 | 0.030 | 0.273 | 0.042 | 0.071 | 0.058 | 0.148 | 0.123 |
|  | 75 | 0.364 | 0.479 | 0.391 | 0.333 | 0.204 | 0.454 | 0.383 | 0.353 | 0.519 | 0.521 | 0.428 |
| L-DOPA | 25 | 0.060 | -0.025 | -0.073 | -0.011 | -0.098 | 0.026 | -0.086 | -0.047 | 0.019 | 0.022 | 0.008 |
|  | 50 | 0.272 | 0.165 | 0.130 | 0.178 | 0.070 | 0.278 | 0.098 | 0.084 | 0.190 | 0.127 | 0.089 |
|  | 75 | 0.574 | 0.517 | 0.383 | 0.390 | 0.291 | 0.367 | 0.346 | 0.374 | 0.598 | 0.508 | 0.492 |

**Figure S1.**
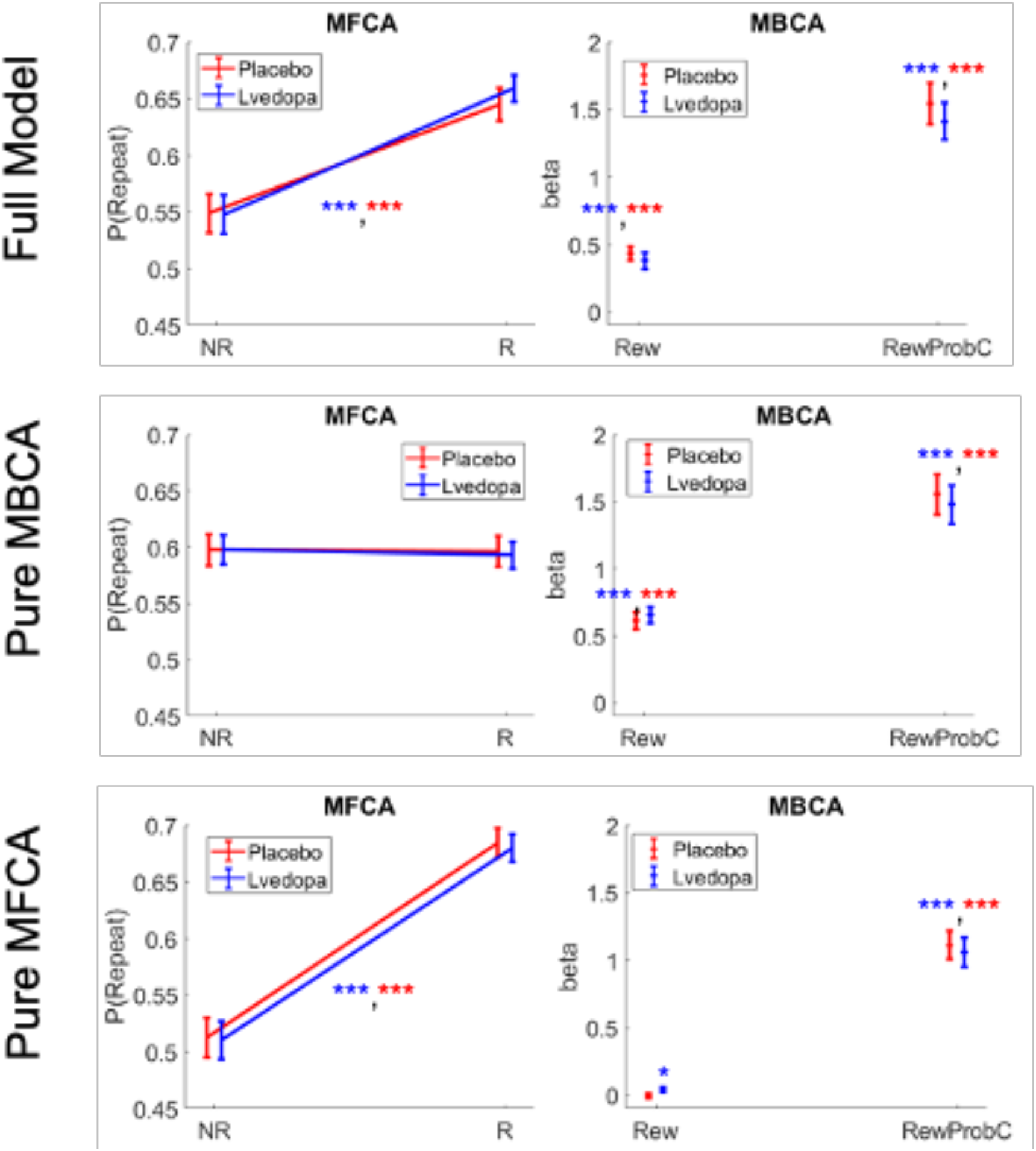
Simulations for standard trials based on the full model and sub-models. NR=no reward, R=reward. Rew=reward at the common destination, RewProBC=Reward Probability at the common destination.

**Figure S2.**
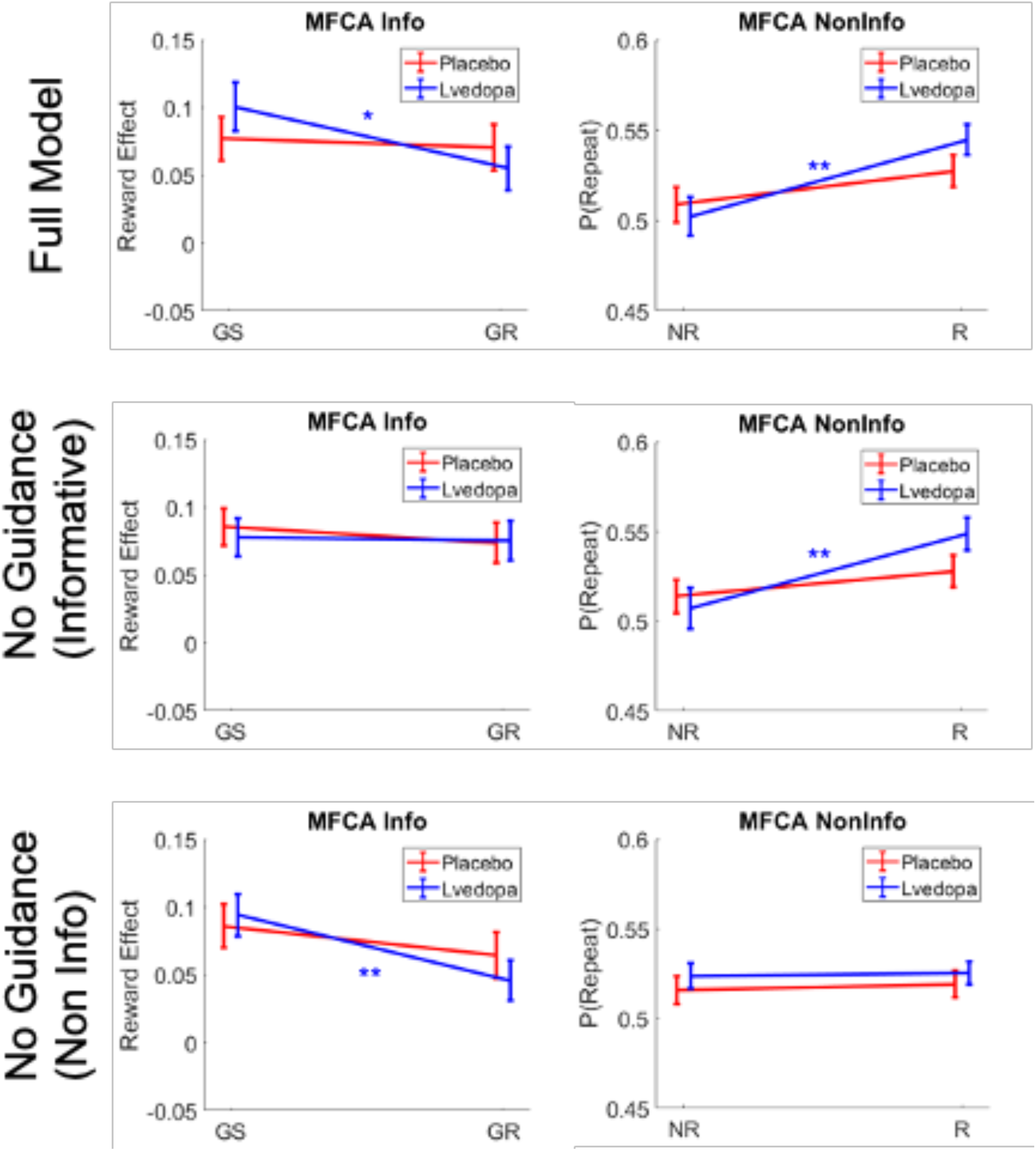
Simulations for uncertainty trials based on the full model and sub-models. GS=Ghost-selected, GR=Ghost-rejected.

**Figure S3.**
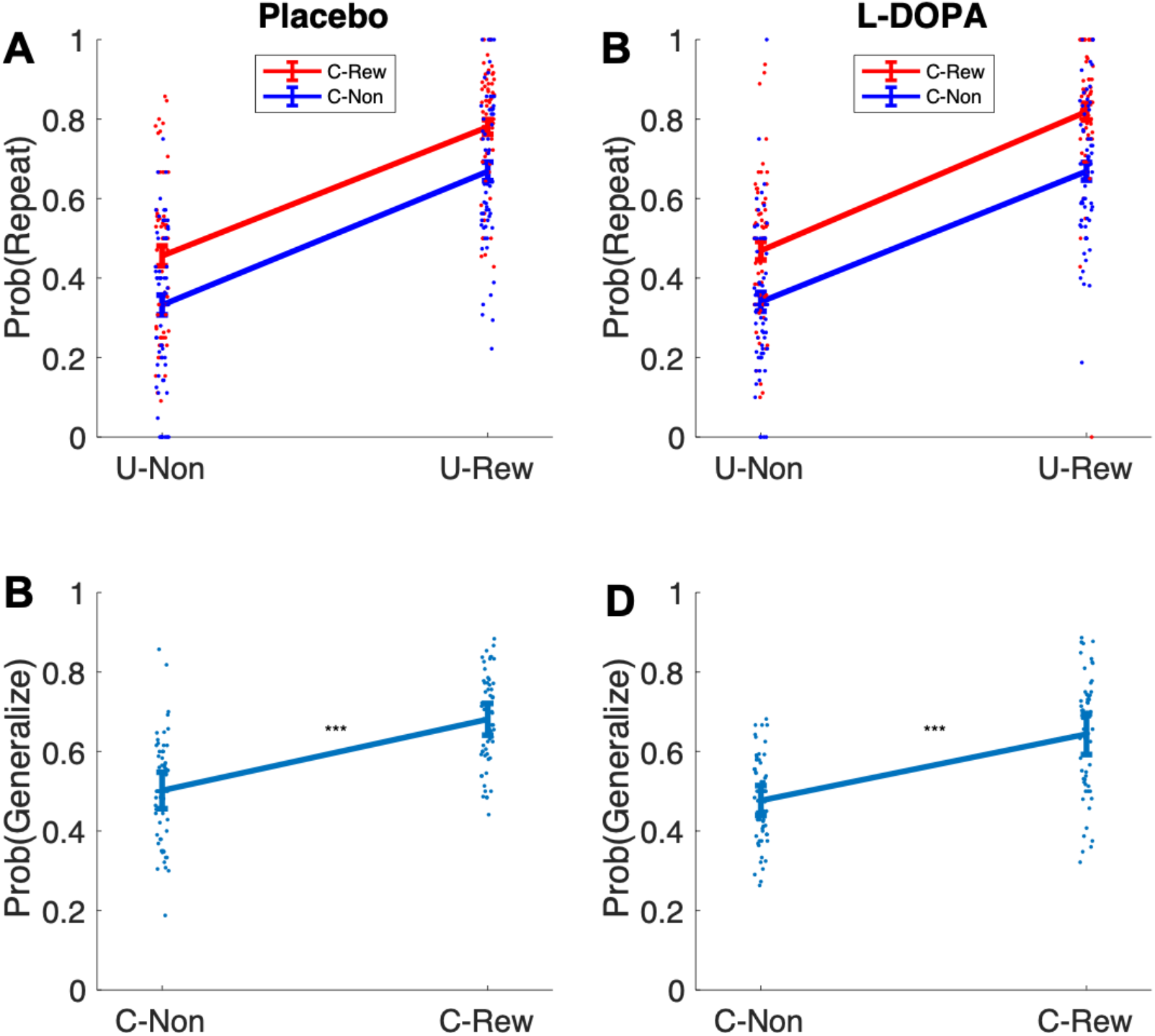
Empirical probabilities of model-agnostic MF (A & B) and MB (C & D) choice contribution under placebo and levodopa (L-DOPA). U-Non=no reward at unique destination, U-Rew= reward at unique destination, C-Non=no reward at common destination, C-Rew= reward at common destination.

**Figure S4.**
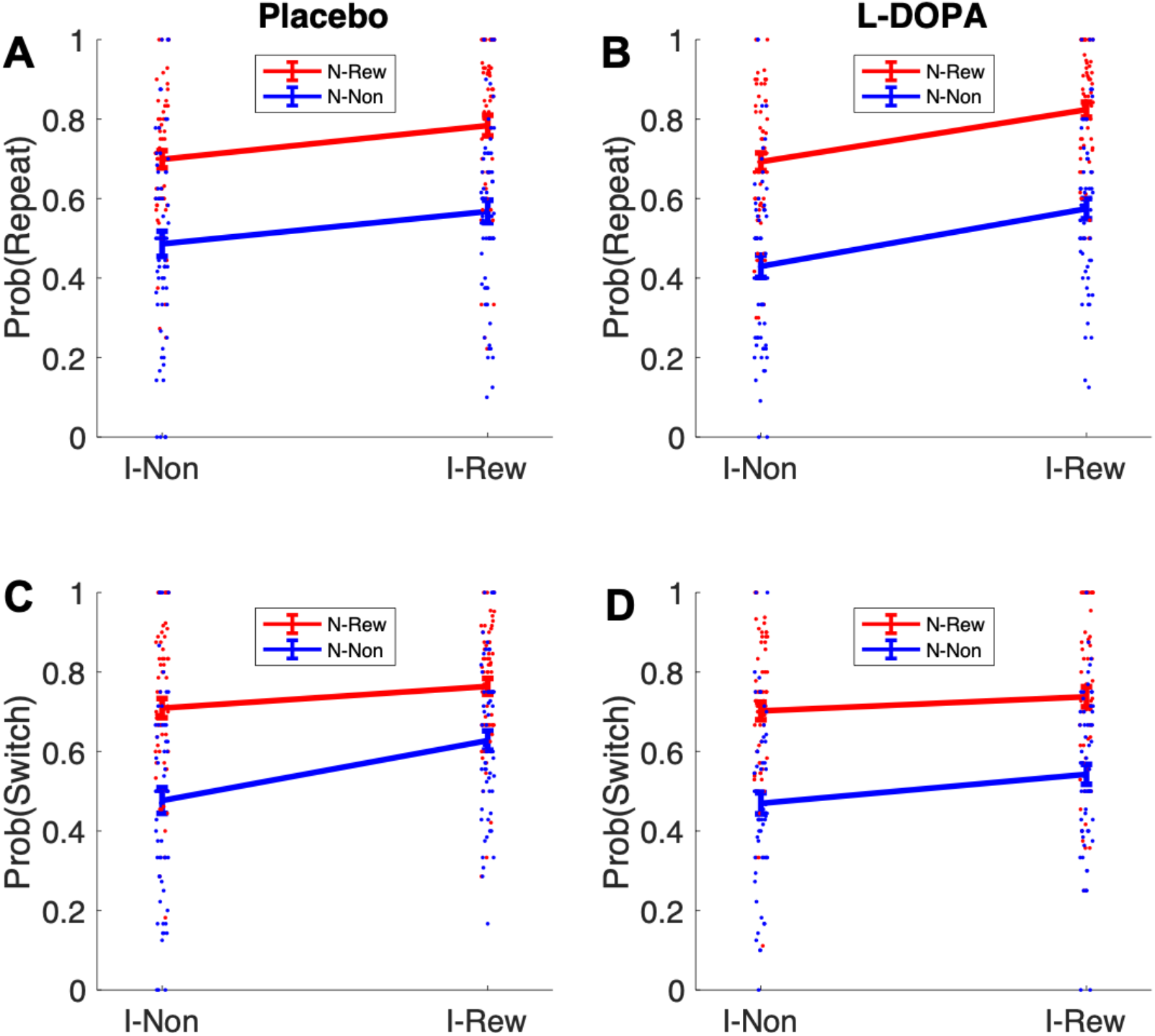
Retrospective MB inference using the informative destination based on repeat and switch signatures after uncertainty trials. I-Non=no reward at informative destination, I-Rew= reward at informative destination, N-Non=no reward at non-informative destination, N-Rew= reward at non-informative destination.

**Figure S5.**
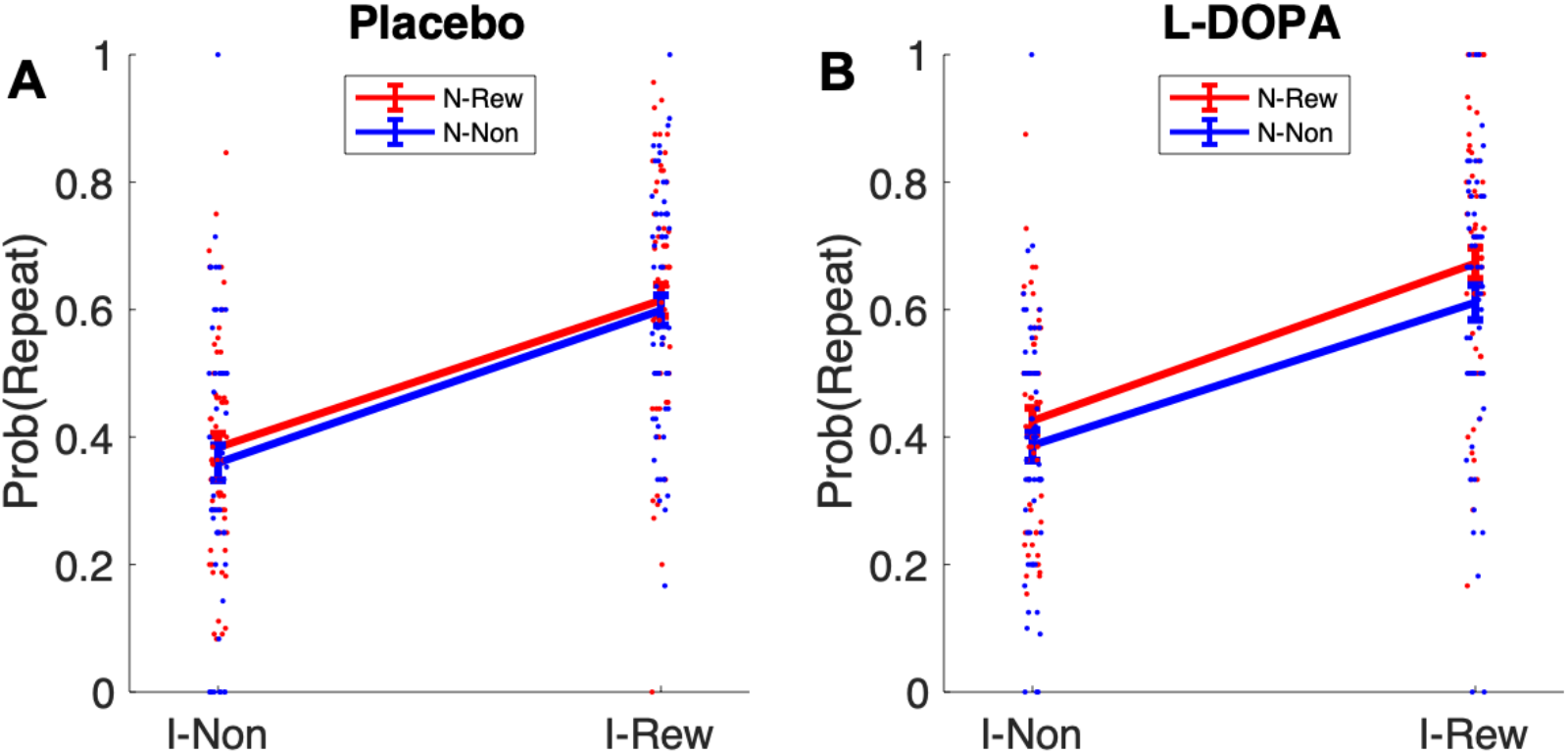
Retrospective MB inference using the non-informative destination based on choice repetition in “clash” trials n+1 following an uncertainty trial-n. I-Non=no reward at informative destination, I-Rew= reward at informative destination, N-Non=no reward at non-informative destination, N-Rew= reward at non-informative destination.

**Figure S6.**
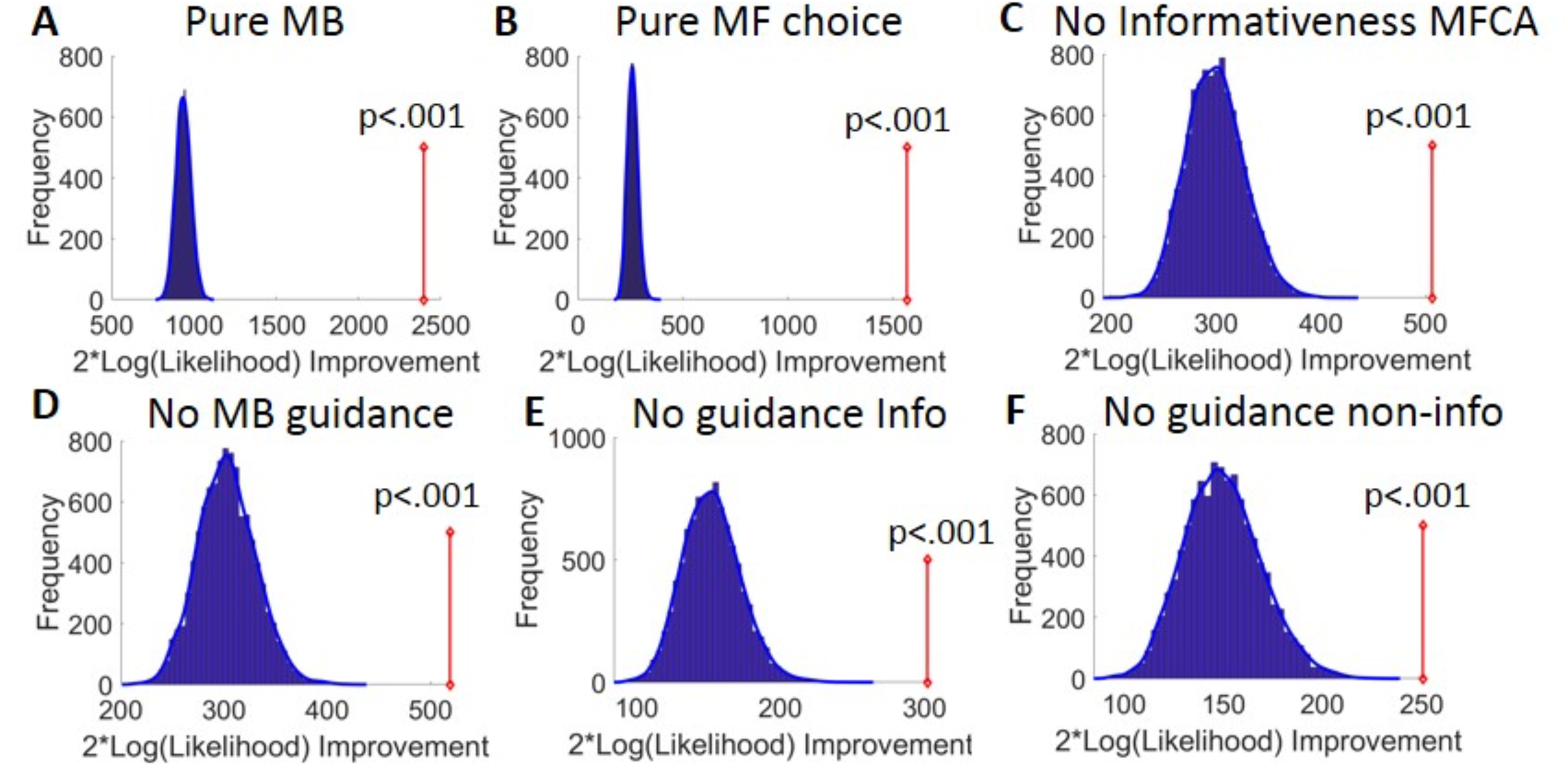
Model-comparison results. A) Results of the bootstrap-GLRT model-comparison for the pure MB sub-model. The blue bars show the histogram of the group twice log-likelihood improvement (model vs. sub-model) for synthetic data simulated using the sub-model (10000 simulations). The blue line displays the smoothed null distribution (using Matlab’s “ksdensity”). The red line shows the empirical group twice log-likelihood improvement. p-value reflect the proportion of 10000 simulations that yielded an improvement in likelihood that was at least as large as the empirical improvement. B-E) Same as (A), but for the pure MF choice, the no informativeness effects on MFCA, the no MB-guidance for MFCA, the no MB-guidance for the informative destination and the no-MB guidance for the non-informative destination sub models.

**Figure S7.**
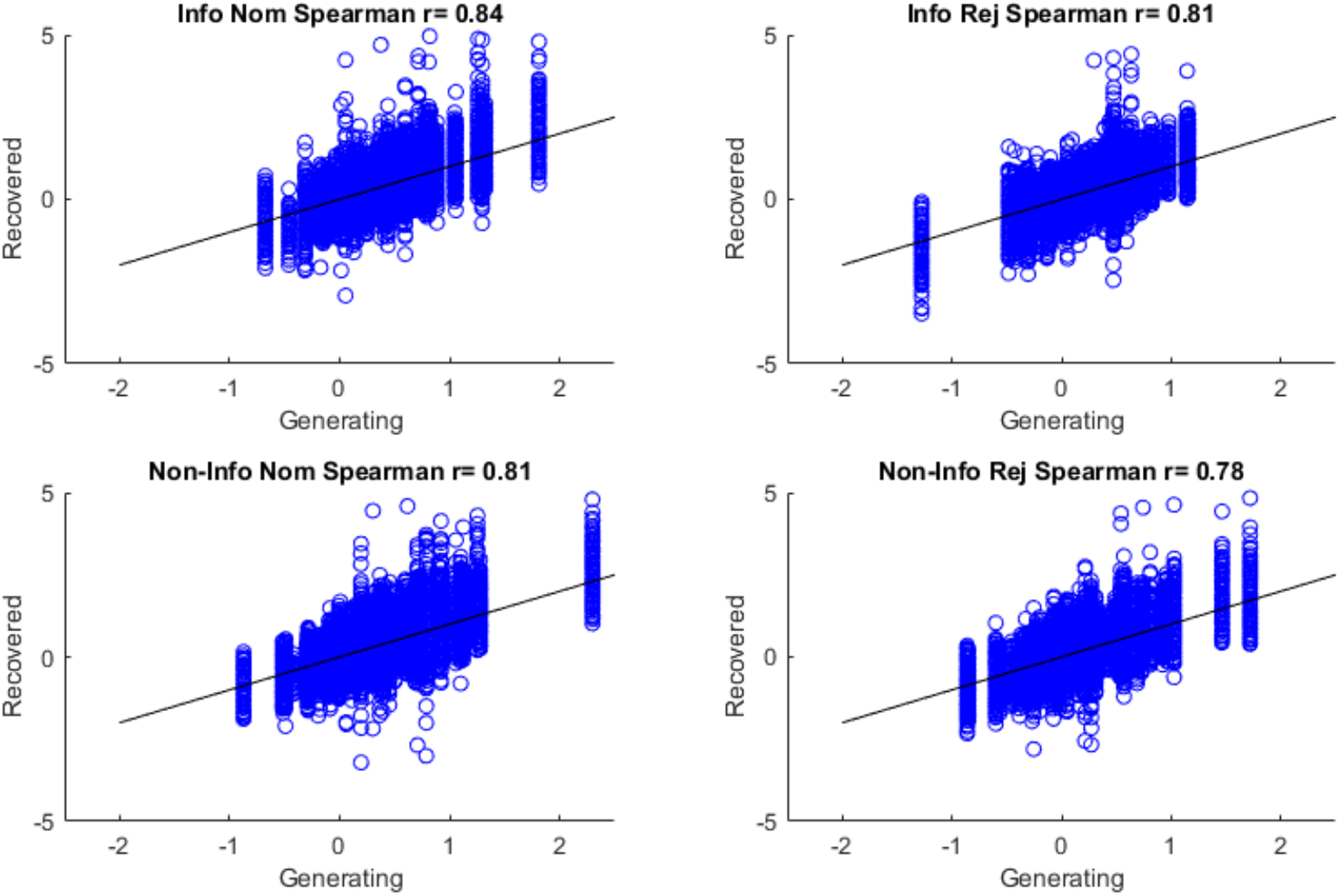
Parameter recoverability. For each of the 2*62 full model parameter-combinations 1000 synthetic (simulated) datasets were created by simulating the full model on experimental sessions as in the true experiment. Then the full model was fit to each of these generated datasets. For each MFCA parameter (info/non-info × nom/rej), we plot the recovered against the generating parameters (and impose black diagonals where “recovered = generating”).

**Figure S8.**
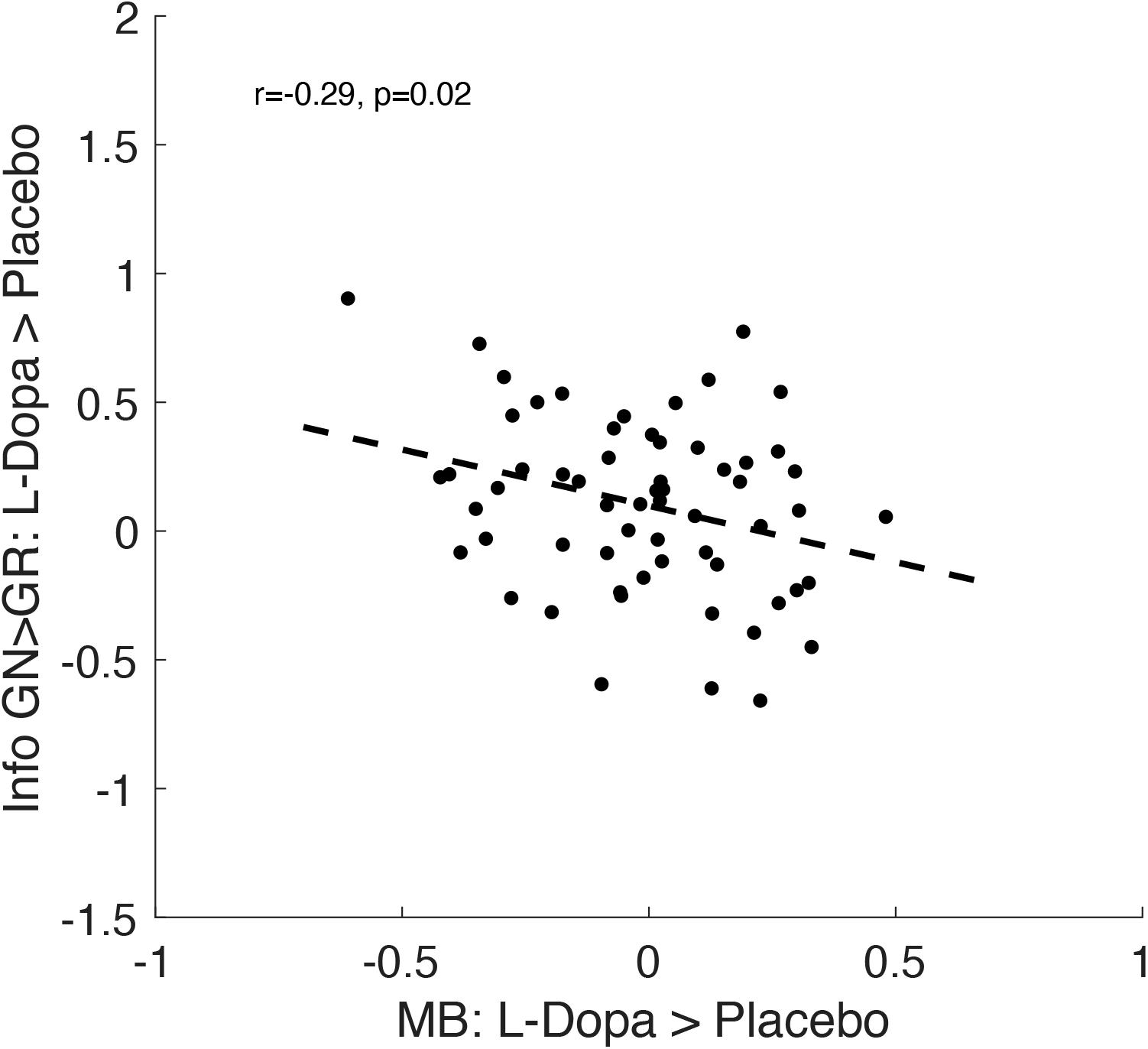
When using a model-agnostic measure of MB choice (probability to generalize after reward minus no-reward) and of preferential MFCA at the informative destination (repeat or ghost-nominate minus switch or ghost-rejected), dopamine dependent differences (levodopa minus placebo) in those measures were correlated negatively (r=-.29, p=.021) mirroring the finding as reported on parameters from the computational model in the main text.

